# Spatial tumor immune heterogeneity facilitates subtype co-existence and therapy response via AP1 dichotomy in pancreatic cancer

**DOI:** 10.1101/2023.10.30.563552

**Authors:** Lukas Klein, Mengyu Tu, Niklas Krebs, Laura Urbach, Daniela Grimm, Muhammad Umair Latif, Frederike Penz, Nathan Chan, Kazeera Aliar, Foram Vyas, Uday Kishore, Elisabeth Hessmann, Andreas Trumpp, Elisa Espinet, Argyris Papantonis, Rama Khokha, Volker Ellenrieder, Barbara T. Grünwald, Shiv K. Singh

## Abstract

Pancreatic ductal adenocarcinoma (PDAC) displays a high degree of spatial subtype heterogeneity. This intratumoral co-existence of classical and basal-like programs is evident in multi-scale transcriptomic and spatial analyses of resected, advanced-stage and chemotherapy-treated specimens and reciprocally linked to a diverse stromal immune microenvironment as well as worse clinical outcome. However, the underlying mechanisms of intratumoral subtype heterogeneity remain largely unclear. Here, by combining preclinical models, multi-center clinical, bulk and compartment-specific transcriptomic, proteomic, and bioimaging data from human specimens, we identified an interplay between neoplastic intrinsic AP1 transcription factor dichotomy and extrinsic CD68^+^ macrophages as a driver of intratumoral subtype co-existence along with an immunosuppressive tumor microenvironment with T cell exclusion. Our ATAC-, ChIP-, and RNA-seq analyses revealed that JUNB/AP1- and HDAC-mediated epigenetic programs repress pro-inflammatory immune signatures in tumor cells, antagonizing cJUN/AP1 signaling to favor a therapy-responsive classical neoplastic identity. Through the tumor microenvironment, this dichotomous regulation was further amplified via regional macrophage populations. Moreover, CD68^+^/TNF-α^+^ cells associated with a reactive phenotype and reduced CD8^+^ T cell infiltration in human PDAC tumors. Consequently, combined anti-TNF-α immunotherapy and chemotherapy resulted in reduced macrophage counts and promoted CD3^+^/CD8^+^ T cell infiltration in basal-like PDAC, leading to improved survival in preclinical murine models. We conclude that tumor cell intrinsic epigenetic programs, together with extrinsic microenvironmental cues, facilitate intratumoral subtype heterogeneity and disease progression.

## Introduction

The molecular heterogeneity in neoplastic and stromal immune cells renders PDAC prognosis dismal and therapy challenging. PDAC has become the third leading cause of cancer-related death with a 5-year survival rate of 11%^1^. Presently, gemcitabine/nab-paclitaxel and modified FOLFIRINOX are the main therapeutic options in PDAC, though therapy resistance and local as well as distal recurrences are common^2,3^. Transcriptome analyses identified two clinically relevant PDAC subtypes; the basal-like (BL) or squamous subtype is linked to therapy resistance and worse patient outcome, whereas the classical (CLA) subtype shows better clinical outcome^4,5^.

Subtype-based screening of a small cohort of PDAC patients has shown potential prognostic as well as predictive benefit^6–8^, and hence, a number of current clinical studies (e.g. NCT05314998) are designed to translate these subtypes into the clinical setting^9^. However, it has become clear that the CLA and BL subtypes are not discrete states of individual PDAC tumors, but rather co-exist and contribute to significant intratumoral heterogeneity that is poorly understood. Multi-scale transcriptomic and imaging-based profiling has revealed a widespread co-existence of CLA and BL subtypes within PDAC patient tumors, as well as hybrid/co-expressor states that are implicated as transition mechanism between the subtype extremes^10–15^, emphasizing the complex tumor cell plasticity. Moreover, the extent of subtype co-existence increases in advanced disease, especially in metastatic samples^13,16,17^. This significantly associates with poorer overall prognosis, making precision-based therapies for PDAC patients a major clinical challenge^13,16,17^. Thus, understanding the underlying mechanisms of intratumoral subtype plasticity may improve subtype-based prediction and therapeutic response for PDAC patients.

While the precise mechanisms that drive this spatial plasticity in PDAC are yet unclear, it appears that both tumor cell intrinsic and extrinsic factors play a role. A major extrinsic driver of malignant cell phenotypic plasticity in PDAC is the regional tumor immune microenvironment (TiME), which enables the acquisition of therapy resistance and aggressive behavior^18,19^. A recent study showed that TGF-β can promote BL subtype specificity and therapy resistance by regulating neoplastic cell-intrinsic transcriptional programs^20^. Furthermore, TNF-α- and IFN-α/γ-mediated signaling events can drive the BL subtype-specific transcriptional program and promote PDAC aggressiveness^20–22^. Notably, this transcription-based subtype plasticity is independent of genetic alterations^20,21^. It is currently unknown whether plasticity imposed by microenvironmental cues or other factors is responsible for intratumoral subtype heterogeneity and how such factors might promote a specific cell-type identity and response to therapy.

PDAC neoplastic cells indeed show a remarkable capacity to change their phenotypic identity through transcriptional and epigenetic regulation. Lineage-specific transcription factors (TFs) and epigenetic co-regulators are considered a key hallmark of PDAC subtype specificity and disease progression^4,21,23–26^. For instance, MYC, TP63, and AP1 are crucial TFs in squamous/BL and inflammatory PDAC subtypes, while TFs such as GATA6 drive the therapy responsive CLA neoplastic identity^7,21,27–29^. Notably, the AP1 inflammatory TF family drives a strong response to external stimuli such as growth factors and cytokines and regulates key cellular processes including differentiation and growth, also in the context of tumor biology^30,31^. The JUN/AP1 factors furthermore exhibit substantial heterogeneity in gene expression in the distinct PDAC subtypes. For instance, while cJUN/AP1^high^ PDAC patients exhibit resistance to gemcitabine/nab-paclitaxel chemotherapy, earlier relapse, and a BL phenotypic state^21,32^, JUNB/AP1 expression is linked to low-grade/CLA-like PDAC^21,23^. This study thus addresses the question whether and how AP1 heterogeneity could regulate intratumoral subtype plasticity, inflammatory programs, and therapy response in PDAC.

We report a novel spatially regulated dichotomy in the AP1 transcriptional programs (JUNB vs. cJUN) that shapes regional subtype identity in PDAC via both tumor cell intrinsic and extrinsic mechanisms. We show that JUNB/AP1- and HDAC-mediated epigenetic and transcriptional networks restrict macrophages infiltration in the TME. These spatial CD68^+^/TNF-α^+^ macrophages promote intratumoral subtype co-existence by destabilizing CLA neoplastic identity and promoting a BL state. Mechanistically, the loss of JUNB-mediated repressive functions is linked to TNF-α signaling, CLA-to-BL neoplastic transition, and poorer outcome in PDAC patients. Notably, macrophage-derived TNF-α^high^ expression marked a reactive stroma with low CD3^+^/CD8^+^ T cell counts in PDAC patient tumors. Combined anti-TNF-α and standard chemotherapy reduced CD68^+^/TNF-α^+^ macrophages and restored CD3^+^/CD8^+^ T cell infiltration, improving the overall outcome in preclinical models. These molecular insights may help define new therapeutic vulnerabilities and subtype-based precision therapy strategies cognizant of intratumoral subtype co-existence in PDAC.

## Results

### JUNB associates with GATA6^high^ CLA identity in PDAC patients

Given their ability to integrate extrinsic inflammatory signals and intrinsic transcriptional programs, AP1 TFs JUNB and cJUN warrant special attention in the context of intratumoral subtype plasticity in PDAC. We first focused on JUNB, since we and others have indicated its involvement in low-grade/CLA-like PDAC identity^21,23^. Using our JUNB ChIP-seq and ATAC-seq analysis of CLA cell lines, we noted enrichment of pathways related to ‘cell adhesion’ and ‘developmental morphogenesis’ (**Fig. 1a**), supporting the notion that JUNB could promote epithelial features and/or early pancreatic differentiation. We therefore queried several publicly available cohorts of PDAC patient transcriptomes^4,33,34^ (n=582) to investigate an association between JUNB and the *bona fide* CLA marker GATA6 (**Extended Data Fig. 1a-c**) and noted a trend towards positive association with GATA6 in the TCGA dataset (**Extended Data Fig. 1a**). As all three studies relied on bulk expression, stromal JUNB and stromal GATA6 may influence these associations. Hence, we analyzed tissues of 105 PDAC patients where JUNB and GATA6 protein expression in malignant neoplastic epithelial cells was carefully annotated (**Extended Data Fig. 1d**). Interestingly, JUNB expression was not uniform across each tumor, but rather displayed varying degrees of intratumoral spatial heterogeneity. Since multiple TMA cores were available for each patient, we were able to quantify regional expression differences and observed intratumoral co-existence of JUNB^high^ and JUNB^low^ regions in 32.4% of patients (**Extended Data Fig. 1e**). We thus tested association of JUNB with the CLA subtype markers GATA6 and E-cadherin (ECAD) both at the patient level as well as in individual TMA cores. JUNB^high^ vs. JUNB^low^ patient samples showed significantly elevated GATA6 expression (**Fig. 1b-d**). Importantly, patient-paired analysis showed that GATA6 expression was higher in JUNB^high^ than JUNB^low^ regions within the same patient (**Fig. 1e**). The epithelial adhesion molecule ECAD showed an analogous elevated expression in JUNB^high^ patients and samples, though it was only significant at the regional level (**Extended Data Fig. 1f-h**). Accordingly, we observed an overall positive correlation of JUNB and GATA6 expression within patients and within individual TMA cores (**Extended Data Fig. 1f**).

**Figure 1.**
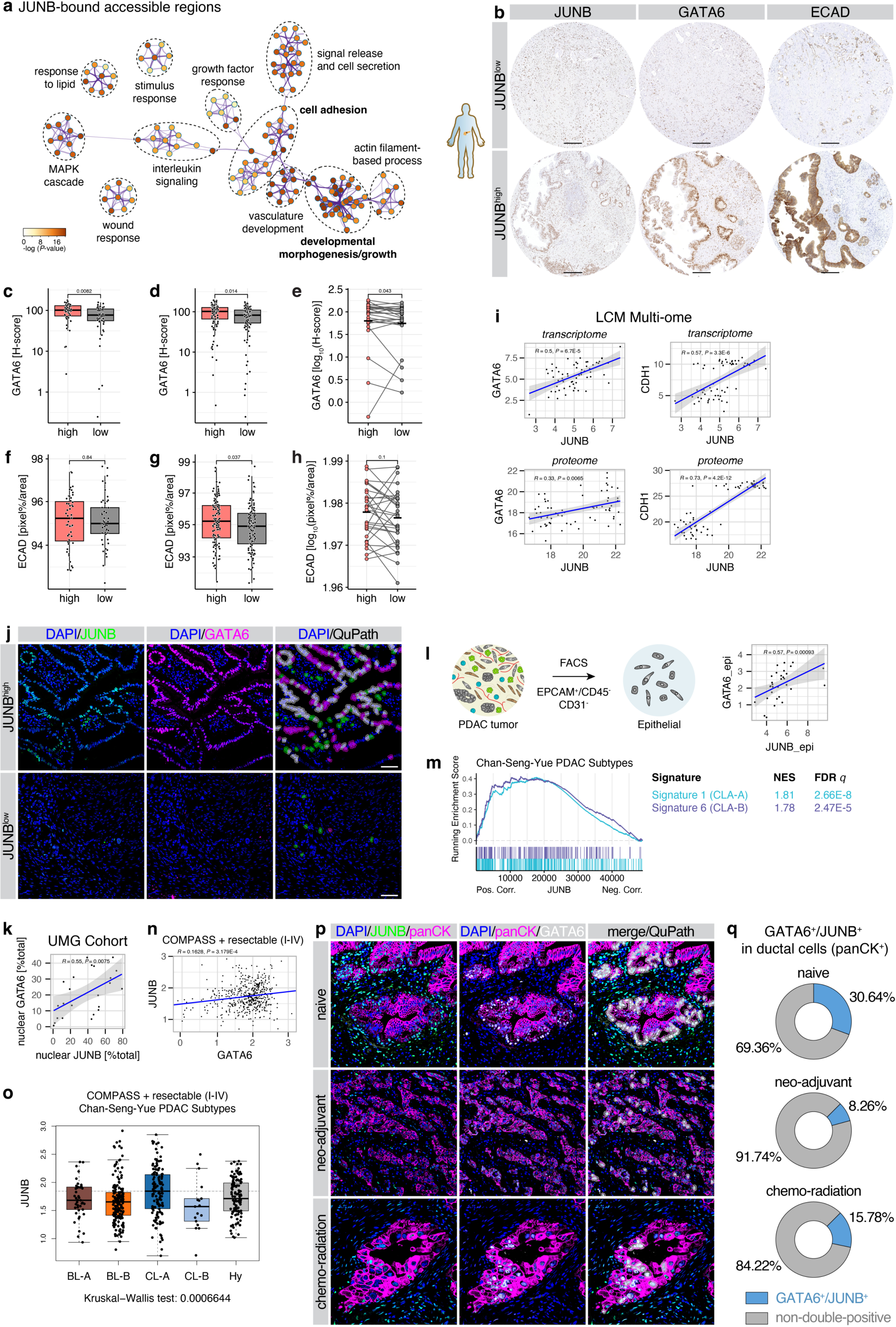
Neoplastic JUNB expression associates with GATA6 in PDAC patients. **a**, Meta-analysis of enriched pathways in regions accessible in CLA PDAC cell lines CAPAN1 and CAPAN2 (by ATAC-seq) and bound in CAPAN1 by JUNB (by ChIP-seq). Node color indicates significance of enrichment, link width the number of overlapping genes between gene sets. **b**-**h**, IHC analysis in 105 PDAC patients for epithelial JUNB expression. **b**, IHC for JUNB, GATA6 and ECAD in cores classified as JUNB^low^ and JUNB^high^. Scale bar 200 μm. **c**-**h**, Quantification of **b**, for GATA6 (**c**-**e**) and ECAD (**f**-**h**) in JUNB^low^ and JUNB^high^ expression per patient (**c**,**f**), per TMA core across all patients (**d**,**g**) and in heterogeneous patients showing matched levels in JUNB^low^ and JUNB^high^ cores (**e**,**h**). **i**, Correlation of JUNB and GATA6 (left), as well as JUNB and CDH1 (right) in laser-capture microdissection-enriched human PDAC tumor epithelia and stroma. Top, RNA-seq expression (transcriptome, n=29 patients), bottom mass-spectrometry quantification (proteome, n=32 patients). Linear regression with 95% CI, as well as Spearman’s *R* and associated *P* value. **j**, IF for JUNB and GATA6 in resection tissue of PDAC patients at representative JUNB^high^ and JUNB^low^ regions, with overlayed cell classification by QuPath. Green: nuclear JUNB^+^, magenta: nuclear GATA6^+^, white: JUNB-GATA6 double-positive. Scale bar 50 μm. **k**, Quantification of **j** for per-patient average nuclear GATA6^+^ cells and nuclear JUNB^+^ cells relative to the total number of cells, plotted as in **i**. n=23. **l**, Epithelial-specific transcriptional RNA-seq profiles of resected PDAC patients were generated by fluorescence-activated cell sorting of EPCAM^+^/CD45^−^/CD31^−^ cells. Correlation analysis for epithelial-specific JUNB and GATA6, plotted as in **i**. n=31. **m**, Gene set enrichment analysis for Chan-Seng-Yue PDAC subtypes^12^ in genes correlating with JUNB in epithelial compartment-sorted transcriptomes of **l**. Normalized enrichment score (NES) and FDR *q* value are indicated. **n**, Correlation analysis for JUNB and GATA6 in LCM-enriched epithelia of patients of the COMPASS trial (stage I-IV). Linear regression, as well as Spearman’s *R* and associated *P* value. n=486. **o**, JUNB expression in the dataset as in **n**, classified for the Chan-Seng-Yue PDAC subtypes. Kruskal-Wallis test. **p**, IF for JUNB, pan-cytokeratin (panCK), and GATA6, in tissue of treatment-naive, neo-adjuvant-treated, and chemo-radiation-treated PDAC patients. Overlayed cell classification by QuPath for panCK-JUNB-GATA6 triple-positive cells shown in white. Scale bar 50 μm. **q**, Quantification of **p** for average percentage of ductal (cytoplasmic panCK^+^ cells) that are additionally double-positive for nuclear GATA6 and nuclear JUNB (blue) or not (grey). Naive, n=4; neo-adjuvant, n=4; chemo-radiation, n=3.

Complementary to the IHC analysis of epithelial JUNB expression, we queried transcriptome and proteome data of human PDAC specimens that were epithelium- and stroma-enriched by laser-capture microdissection^35^. Again, a strong correlation of JUNB with GATA6, as well as with ECAD (*CDH1*) was evident (**Fig. 1i**). Furthermore, dual IF staining of JUNB and GATA6 in a small cohort of 23 patients (**Fig. 1j**) showed a matching strong correlation of nuclear JUNB and GATA6 expression in epithelial cells (**Fig. 1k**). In addition, flow cytometry-sorted epithelial-specific (EPCAM^+^/CD45^−^/CD31^−^) transcriptomes of 31 PDAC patients^22^ further confirmed a positive link between JUNB and GATA6 expression (**Fig. 1l**). Finally, we observed an enrichment of CLA-A and CLA-B subtype signatures of Chan-Seng-Yue *et al.*^12^ with JUNB expression in the epithelial compartment of patients (**Fig. 1m**), in line with the association with the CLA subtype marker GATA6. Given this positive association between epithelial JUNB and GATA6 in multiple cohorts of resected stage I-II PDAC specimens, we proceeded to investigate whether this association was maintained at later stages. We queried the epithelial-enriched RNA expression dataset from the COMPASS trial^7^, which includes data of LCM-enriched epithelia of early and advanced stage patients (stage I-IV). Here, JUNB again correlated with GATA6 (**Fig. 1n**), and patients classified as CLA-A^12^ showed the highest expression of JUNB (**Fig. 1o**), in line with the epithelial-specific transcriptome data (**Fig. 1m**). PDAC has been described to show a remarkable level of plasticity in response to extrinsic signals, via microenvironmental stimuli as well as treatment^16,19,21,36,37^. Hence, we assessed the expression of neoplastic JUNB and GATA6 in a set of treatment-naive, neoadjuvant chemotherapy-treated, and chemo-radiation-treated PDAC specimens (**Fig. 1p**, **Extended Data Fig. 1g**). In all treated and non-treated samples, most GATA6^+^ neoplastic (panCK^+^) cells also expressed JUNB (77.5%). Additionally, GATA6/JUNB double-positive ductal cells were less abundant upon treatment, especially in neoadjuvant samples (**Fig. 1l, Extended Data Fig. 1h**). Despite the limited number of samples for this analysis, the data further supports the association of JUNB^+^/GATA6^+^ states with the CLA, possibly chemoresponsive identity, and underlines its instability upon therapy.

### JUNB-mediated transcriptional repression affects the CLA phenotype, inflammation, and clinical outcome

As higher JUNB levels were linked to a GATA6-positive, CLA-like phenotype in PDAC neoplastic cells (**Fig. 1**), we investigated the underlying transcriptional mechanism. We utilized our ChIP-seq data for JUNB (as in **Fig. 1a**), together with publicly available H3K27ac^23^ data to determine potential direct regulatory effects of JUNB on lineage TFs of the CLA subtype. JUNB binds not only on itself (**Fig. 2a**), but crucially also on a potential downstream enhancer of GATA6 (**Fig. 2b**). Other CLA-associated factors such as HNF1B and FOXA1 are also bound directly by JUNB in CLA cells at intronic and promoter regions, respectively (**Fig. 2c,d**). Both JUNB binding (**Fig. 2e**) and H3K27ac occupancy (**Fig. 2f**) were validated via ChIP-qPCR in CLA cells, which showed the strongest binding of JUNB at the GATA6 locus. Next, regulatory effects of JUNB were investigated in a global approach by integrating JUNB-bound regions by ChIP-seq with differential expression upon silencing of JUNB (siJUNB) compared to negative control siRNA (siCtrl; **Extended Data Fig. 2a**). Genes directly bound by JUNB showed a higher fold change in gene expression than all genes (**Fig. 2g**). As further illustrated in **Fig. 2h**, JUNB-bound genes were preferentially upregulated upon silencing, suggesting a direct repression by JUNB. Gene ontology analysis of genes directly bound by JUNB indicated that pathways involved in cell migration/stemness, inflammatory signaling, as well as histone deacetylase (HDAC) targets were enriched upon JUNB silencing in PDAC cells (**Fig. 2i**). Notably, the ‘JUNB repression signature’ contained major BL-specific driver genes, such as *CD9*, *MYC*, *TP63,* and *cJUN* (**Supplementary Table 3**), indicating a direct tumor cell-intrinsic repression of BL features. In accordance, JUNB silencing in established (CAPAN2) as well as in PDX-derived (JUNB^high^ GCDX62) CLA cell lines led to a more invasive state (**Extended Data Fig. 2b-g**), which is a characteristic of BL PDAC cells.

**Figure 2.**
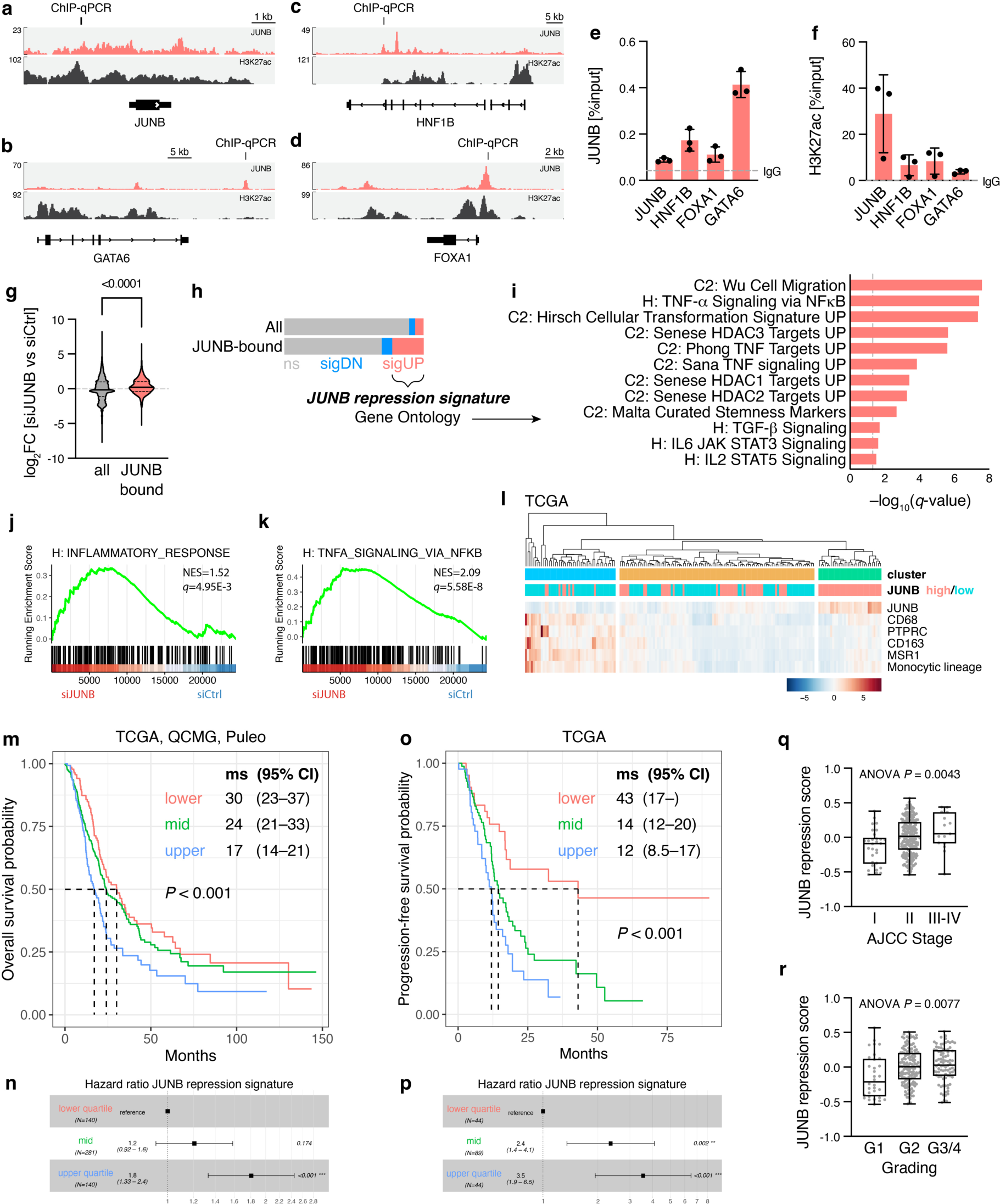
Prognostic relevance of JUNB-repressed inflammatory signaling. **a**-**d**, Coverage of previously published^21^ JUNB ChIP-seq data in CAPAN1, as well as publicly available H3K27ac^23^ data, for loci of JUNB (**a**), GATA6 (**b**), HNF1B (**c**), and FOXA1 (**d**). ChIP-qPCR validation regions are indicated. **e**,**f**, ChIP-qPCR for regions indicated in **a**-**d**, showing signal relative to input for JUNB (**e**) and H3K27ac (**f**) pulldown with mean ± s.d. and average IgG isotype control. n=3. **g**-**i**, Integration of RNA-seq data performed after JUNB silencing (siJUNB; n=3) or control siRNA (siCtrl; n=2) in CAPAN1, with ChIP-seq for JUNB. **g**, Violin plot of log_2_ fold change (FC) in siJUNB RNA-seq data for all (n=36.740) or JUNB-bound (n=698) genes. Median and quartiles are indicated. Student’s t-test with Welch’s correction. **h**, As in **g**, showing the number of genes that display a significant upregulation (sigUP) or downregulation (sigDN), or no significant change (ns). **i**, Gene ontology analysis of significantly upregulated, JUNB-bound genes following JUNB silencing with –log_10_(*q*-value) indicated. Hallmark (H) and curated (C2) signature collections of the Molecular Signature Database (MSigDB) are shown. **j**,**k**, Gene set enrichment analysis plots for “inflammatory response” (**j**) and “TNF-α signaling via NFκB” (**k**) Hallmark signatures of the MSigDB for siJUNB versus siCtrl in CAPAN1 cells. Normalized enrichment score (NES) and FDR *q* value are indicated. **l**, Heatmap of TCGA expression data for JUNB and macrophage markers as well as MCPcounter scores for the monocytic lineage. Cell color indicates z score. JUNB high/low annotation based on top/bottom half of patients for JUNB expression. n=177. **m**,**n**, Overall survival and hazard ratio in TCGA (n=177), Puleo (n=288), and QCMG (n=96) patients stratified by JUNB repression signature (**h**) score. **m**, Kaplan-Meier survival analysis for the lower/upper quartiles (n=140 each) and mid group (n=281) for JUNB repression signature scores. Median survival (ms) with 95% confidence interval (CI). Log-rank test. **n**, Cox proportional hazard groups as in **j**. Hazard ratio (to lower quartile) with 95% CI. *P* values are shown right. **o**,**p**, As in **m**,**n**, for progression-free survival in the TCGA cohort. **q**,**r**, JUNB repression signature scores in AJCC stages (**q**) and pathological grading (**r**) for TCGA and QCMG cohorts combined

Furthermore, we noted a significant enrichment of inflammatory response and multiple TNF-α signaling hallmark signatures (**Fig. 2i-k**), as well as TGF-β signaling and IFN-γ response (**Extended Data Fig. 2h,i**), upon JUNB silencing. This was of particular interest since recent studies have shown that TNF-α signaling pathway is linked to a therapy-induced plasticity and macrophage-driven inflammatory response in PDAC patients^21,36^. We thus investigated a potential association between JUNB and macrophage infiltration (based on marker expression and MCPcounter scores) in PDAC patients. Hierarchical clustering of 177 patients revealed nearly a quarter (26.0%) of macrophage^high^/JUNB^low^ patients (blue cluster), while 18.1% showed high JUNB expression and low macrophages (green cluster) (**Fig. 2l**). Though the clusters are more diverse in the Puleo and QCMG data, JUNB^high^ expression was generally associated with clusters of reduced macrophage signatures (**Extended Data Fig. 2j,k**). We then tested the impact of these JUNB regulatory effects on overall clinical outcomes in PDAC patients. We used gene set variation analysis (GSVA) for the stratification of patients by the ‘JUNB repression signature’. In the combined TCGA, QCMG, and Puleo datasets (total n=582), this revealed a clear overall survival benefit for low expression of the JUNB repression signature (17 vs 30 months in upper vs lower quartile; **Fig. 2m**; hazard ratio 1.8 [95% CI 1.3 – 2.4; **Fig. 2n**]). A particularly stark difference was noted in the progression-free survival rate among the TCGA patients (**Fig. 2o**), with a hazard ratio of 3.5 (95% CI 1.9 – 6.5) for high expression of the signature (**Fig. 2p**). Concordantly, JUNB repression score was lowest with lower AJCC stage (**Fig. 2q**) and lower histological grade (**Fig. 2r**). Together, these data suggest that JUNB-dependent repression of intrinsic drivers of the BL phenotype as well as extrinsic inflammatory drivers such as TNF-α signaling confer reduced macrophage recruitment and improved survival in PDAC patients.

### JUNB antagonizes cJUN and cytokine expression utilizing HDAC1

The cJUN TF, an AP1 family member of JUNB, is a crucial mediator of pro-inflammatory signaling and BL identity in PDAC^21,32^. We aimed to investigate whether JUNB-dependent repression of inflammatory pathways is associated with cJUN signaling. First, we carefully validated the expression of core inflammatory factors in the major inflammatory signatures that were repressed by JUNB (**Fig. 2p,q**, **Extended Data Fig. 2h,i**). Several interleukins (e.g. IL-1α/β, IL-6) and C-X-C/C-C motif chemokines showed an expected upregulation upon silencing of JUNB in RNA-seq (**Fig. 3a**) and qPCR (**Fig. 3b**). Interestingly, cJUN itself and its downstream target CCL2 were upregulated upon JUNB downregulation (**Fig. 3b**). ChIP-seq data of JUNB and H3K27ac showed that both cJUN itself (**Fig. 3c**), as well as the loci for IL-1α/β (**Fig. 3d**) and CXCL9/10/11 (**Fig. 3e**) displayed strong JUNB binding in the absence of H3K27ac, indicating repression.

**Figure 3.**
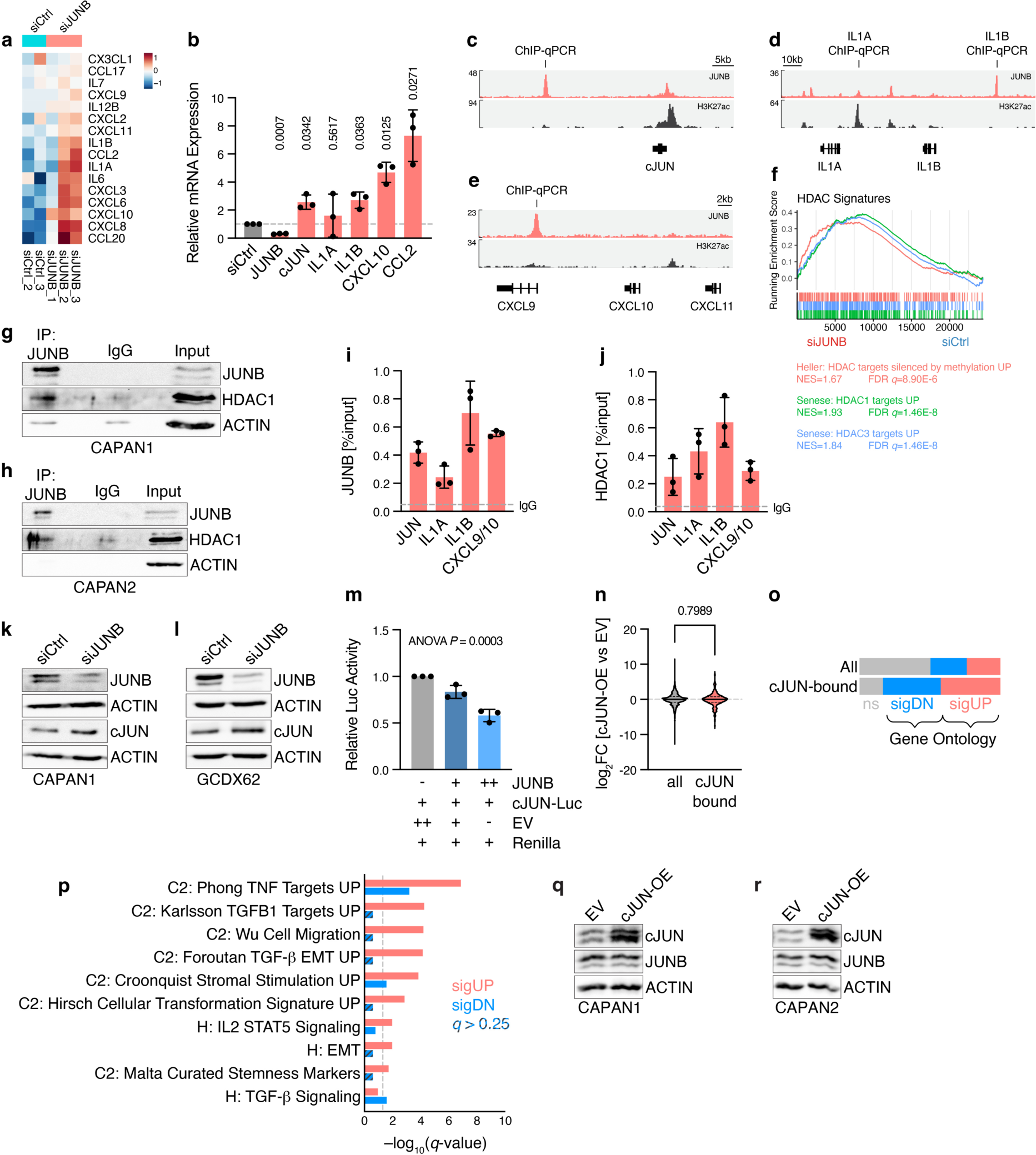
JUNB-HDAC1 complex represses inflammatory signals and cJUN. **a**, Heatmap showing expression of cytokines present in the core enrichment of the gene sets shown in **Figure 2p,q**, and **Extended Data Figure 2h,i**, for JUNB silencing (siJUNB; n=3) versus control siRNA (siCtrl; n=2) in CAPAN1 cells. Cell color indicates z score. **b**, qRT-PCR analysis for indicated target genes in siJUNB conditions (red), normalized to siCtrl (grey), in CAPAN1. Relative mRNA expression with mean ± s.d. shown. n=3. Student’s t-test with Welch’s correction. **c**-**e**, Coverage of JUNB ChIP-seq data in CAPAN1^21^, as well as publicly available H3K27ac^23^ data, for loci of cJUN (**b**), IL1A/B (**c**), and CXCL9/10/11 (**d**). ChIP-qPCR validation regions are indicated. **f**, Gene set enrichment analysis for curated signatures (C2) of the Molecular Signature database (MSigDB) for siJUNB versus siCtrl in CAPAN1 cells. Normalized enrichment score (NES) and FDR *q* value are indicated. **g**,**h**, Immunoblot for JUNB, HDAC1, and β-actin after JUNB pulldown, IgG isotype control or input in CAPAN1 (**g**) and CAPAN2 (**h**). n=3. **i**,**j**, ChIP-qPCR for regions indicated in **c**-**e**, showing signal relative to input for JUNB (**i**) and HDAC1 (**j**) pulldown with mean ± s.d. and average IgG isotype control. n=3. **k**,**l**, Representative immunoblot for JUNB, cJUN, and β-actin in CAPAN2 (**k**) and GCDX62 (**l**) after siJUNB or siCtrl. n=3. **m**, Dual-luciferase reporter assay for cJUN promoter firefly luciferase constructs in CAPAN2 cells transfected with varying concentrations of JUNB overexpression plasmids (or EV controls), together with *Renilla* luciferase control and firefly luciferase (Luc) reporters. Relative Luc activity to control with mean ± s.d. shown. One-way ANOVA. **n**-**p**, Integration of RNA-seq data performed in three biological replicates for overexpression of cJUN (cJUN-OE) or empty vector (EV) control in GCDX62 with ChIP-seq for cJUN. **n**, Violin plot of log_2_ fold change (FC) in cJUN-OE RNA-seq data for all (n=24.118) or cJUN-bound (n=224) genes. Median and quartiles are indicated. **o**, As in **n**, showing the number of genes that display a significant upregulation (sigUP) or downregulation (sigDN), or no significant change (ns). **p**, Gene ontology analysis of significantly upregulated (red) or downregulated (blue), cJUN-bound genes following cJUN-OE with –log_10_(*q*-value) indicated. Hallmark (H) and curated (C2) signature collections of the MSigDB are shown. Not enriched pathways (*q*-value>0.25) are indicated by striped bars. **q**,**r**, Immunoblot for JUNB, cJUN, and b-actin in CAPAN1 (**q**) and CAPAN2 (**r**) cells with overexpression of cJUN (cJUN-OE) or empty vector (EV) control. n=3.

TFs require additional epigenetic co-regulators to exert their transcriptional regulatory functions. In the JUNB repression signature genes (see **Fig. 2i**), as well as in GSEA in JUNB silencing transcriptome data, HDAC target signatures were upregulated (**Fig. 3f**). Therefore, we hypothesized that HDACs may be involved in JUNB-mediated transcriptional repression of inflammatory BL-specific lineage signatures. To determine whether HDAC1 cooperates with JUNB in transcriptional repression, we investigated protein interaction, which confirmed a direct binding between JUNB and HDAC1 (**Fig. 3g,h**). Importantly, ChIP-qPCR analysis further validated significant binding of both JUNB (**Fig. 3i**) and HDAC1 (**Fig. 3j**) at the repressed loci, which, we suspect, deacetylates and thereby represses these inflammatory genes. Depletion of JUNB in established and PDX-derived CLA cell lines caused upregulation of cJUN, indicating a direct repression of cJUN by JUNB (**Fig. 3k,l**). This was further confirmed by dual luciferase reporter assays for the promoter of cJUN, which showed that JUNB was able to directly downregulate cJUN (**Fig. 3m**), suggesting that cJUN plays an antagonistic role, activating BL subtype-associated inflammatory genes. To determine the molecular and functional differences between cJUN and JUNB in subtype plasticity, we analyzed cJUN-bound regions by ChIP-seq upon cJUN overexpression (cJUN-OE) in JUNB^high^ CLA cell lines (**Fig. 3n and Fig. 3o**). Notably, unlike JUNB, cJUN-bound genes showed no preference for up- or downregulation in RNA-seq data following HA-tagged cJUN-OE compared to empty vector (EV) controls (**Fig. 3n**), underlining their diverging functions in PDAC subtype plasticity. Gene ontology analysis of up- and downregulated genes (**Fig. 3o**) showed that the aggressiveness-associated pathways repressed by JUNB (e.g. ‘cell migration’; **Fig. 2i**) were directly activated by cJUN (**Fig. 3p**). In particular, EMT- and TNF-α-associated gene signatures were enriched among cJUN-bound genes (**Fig. 3p**), indicating that cJUN may attenuate CLA-associated functions by antagonizing JUNB signaling. Interestingly, however, JUNB expression was not altered upon cJUN overexpression in CLA cell lines (**Fig. 3q,r**). We thus hypothesized that cJUN employs other indirect pathways to antagonize JUNB-dependent signaling. Together, these findings indicate that JUNB restricts BL pro-inflammatory programs via HDAC-mediated transcriptional repression.

### Antagonistic roles of AP1 factors determine regional immune recruitment

Detailed gene expression analysis has shown potential immune-modulatory effects of the antagonistic JUNB-cJUN interplay (**Fig. 2**). To investigate the antithetical functions of JUNB and cJUN in the TME, we generated a CLA (JUNB^high^) subtype-derived orthotopic mouse model, where we overexpressed HA-tagged cJUN (cJUN-OE) and then orthotopically implanted empty vector control (EV) or cJUN-OE cells (**Fig. 4a**). First, we confirmed the expected high nuclear cJUN expression levels compared to EV in cJUN-OE PDAC tumors (**Extended Data Fig. 3a,b**). Intriguingly though, not all ductal cells in the HA-cJUN-OE tumors showed cJUN expression (**Extended Data Fig. 3a**). As JUNB attenuated expression of cJUN (**see Fig. 3k-m**), we next assessed whether cJUN^low^ areas in this heterogeneous tumor model exhibited high JUNB expression. This presented a valuable opportunity to investigate spatial effects of the AP1 heterogeneity in PDAC. Using whole slide images of IHC for HA-cJUN (**Fig. 4c**) and IF for JUNB (**Fig. 4d**) in serial sections, we marked and quantified ‘hotspot’ regions of high cJUN and high JUNB expression. This revealed not only reduced HA-cJUN^+^ cells in JUNB hotspots, but JUNB^+^ cells were *vice versa* depleted in cJUN hotspot areas (**Fig. 4e**). Since cJUN-OE did not lead to a direct repression of JUNB expression (**Fig. 3q,r**), we hypothesized that microenvironmental factors may affect JUNB expression and shape PDAC plasticity. Previous studies have reported that microenvironmental factors such as TNF-α or TGF-β can influence subtype specificity^20,21^. Interestingly, we observed a trend towards higher immune infiltrations (**Fig. 4b**), marked by an increase in CD45^+^/CD68^+^ and TNF-α^+^/CD68^+^ macrophages in cJUN-OE CLA-derived tumors (**Extended Data Fig. 3c-e**). Next, we sought to determine whether regional CD68^+^ macrophage infiltrations, particularly surrounding cJUN^+^ hotspot area, might destabilize JUNB expression (**Fig. 4f**). Indeed, the average distance of CD68^+^ cells to cJUN hotspots was lower than to JUNB hotspots (448.5 vs 645.3 μm; **Fig. 4g**). When looking at the distance of each cell to either hotspot, considerably more CD68^+^ cells showed a shorter distance to the cJUN hotspot (**Fig. 4h**). Thus, cJUN^+^ cell clusters appeared to recruit macrophages, which was restricted in JUNB^+^ areas, possibly affecting AP1 heterogeneity through spatial inflammatory cues. Collectively, these results suggest that regional inflammatory macrophages can influence neoplastic stability by shaping AP1 transcriptional programs.

**Figure 4.**
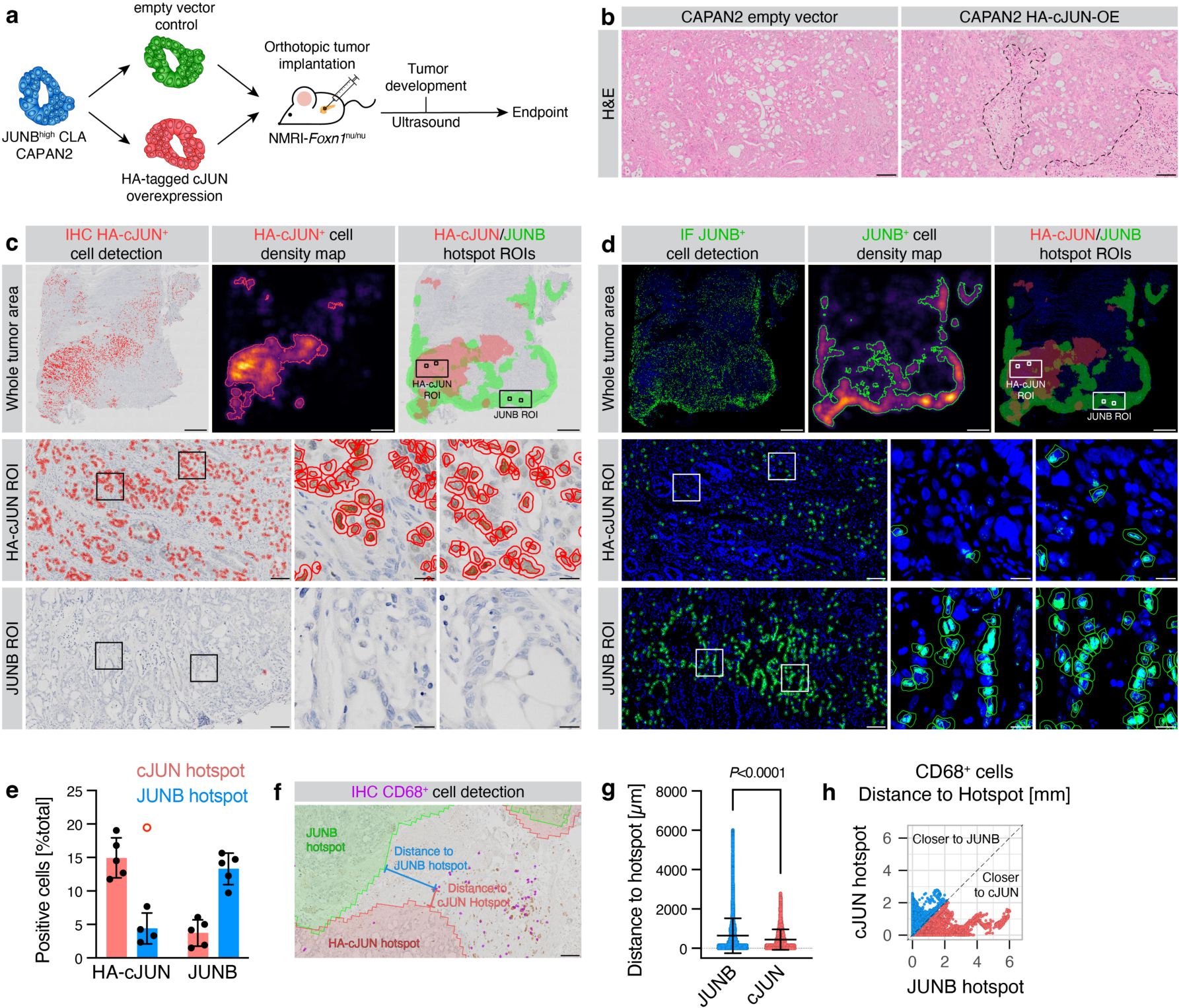
Regional AP1 heterogeneity determines macrophage recruitment. **a**, NMRI-*Foxn1*^nu/nu^ mice were orthotopically transplanted with CAPAN2 cells with stable HA-tagged cJUN overexpression (HA-cJUN-OE) or empty vector (EV) control. **b**, H&E staining of CAPAN2 HA-cJUN-OE and EV tumors. Immune infiltrates are indicated. Scale bar 100 μm. **c**-**h**, QuPath-based analysis of HA-cJUN-OE tumors for HA-cJUN, JUNB, and CD68. **c**,**d**, IHC for the HA tag of cJUN (**c**) or IF for JUNB (**d**) in serial sections. Nuclear positive cell detections (red/green) are indicated. Density maps of positive cells were created and thresholded to derive hotspot regions for HA-cJUN^+^ and JUNB^+^ cells, respectively, in the same tumors. Whole tumor overviews (top) as well as a HA-cJUN (mid) and a JUNB (bottom) hotspot ROIs are shown. **e**, quantification of **c**,**d**, for HA-cJUN^+^ and JUNB^+^ cells relative to the total number of detected cells in cJUN (red) and JUNB (blue) hotspot regions, with mean ± s.d. shown. One outlier is indicated (red circle), which was excluded for mean and s.d. n=5. **f**, IHC for CD68 in HA-cJUN-OE tumors, with CD68^+^ cell detection (purple), as well as JUNB and HA-cJUN hotspots and exemplary 2D distance measurement strategy. **g**, Distance analysis of CD68^+^ cells to JUNB or HA-cJUN hotspots. Scatter plots show each individual cell, with mean ± s.d. Student’s t-test with Welch’s correction. n=14774 CD68^+^ cells from n=5 tumors, with a total of n=1003639 cells analyzed. **h**, As in **g**, showing the shortest distances of each CD68^+^ cell towards both the HA-cJUN and JUNB hotspots.

### TNF-α destabilizes CLA neoplastic identity and shapes local TME heterogeneity

It has been shown that TNF-α or TGF-β can influence subtype specificity^20,21^, yet the impact of inflammatory cell-derived TNF-α on AP1 heterogeneity is unknown. Thus, to determine whether TNF-α destabilizes the CLA subtype (possibly by affecting JUNB signaling) and promotes BL plasticity, we analyzed its impact on transcriptional signatures *in vitro* and *in vivo*. Among genes upregulated in RNA-seq data of CLA PDAC cells upon exogenous TNF-α treatment, we found JUNB, together with a shift in established CLA and BL subtype markers (**Fig. 5a**). To further test this observation *in vivo*, we utilized a CLA-derived orthotopic murine model which was treated for three weeks with exogenous TNF-α. These tumors were then subjected to comprehensive cell type-specific transcriptome analysis. Alignment of bulk RNA-seq of these tumors to human (implanted neoplastic epithelial cells) or murine (host stromal cells) reference genomes and XenofilteR-based removal of the opposite species reads allowed generation of virtually microdissected, compartment-specific transcriptomes (**Fig. 5b**). Within the tumor cell-specific data, GSEA showed a strong enrichment of TNF-α signaling pathways (**Fig. 5c**), confirming that tumor cells reacted to the exogenous treatment. In accordance with the *in vitro* data, TNF-α treatment led to repression of CLA gene signatures *in vivo* (**Fig. 5d**). The stromal population also responded to the TNF-α treatment with induction of TNF-α signaling (**Fig. 5e**), along with a significant remodeling of the stromal immune populations, as determined by MCPcounter (**Fig. 5f**). In particular, cytotoxic T cell, as well as B cell signatures were reduced after TNF-α treatment. Similar to TNF-α treatment, patients with a high expression of the JUNB repression signature genes (see **Fig. 2**) showed a reduction in T cell populations and B cells (**Fig. 5g**). As shown above, the JUNB repression signature was directly associated with TNF-α signaling as well (**Fig. 2i**). Hence, we suspected that JUNB expression might be reduced in response to TNF-α, as a mechanism potentially involved in the indirect cJUN-dependent attenuation of JUNB in the spatial TiME. Indeed, IF staining for JUNB (**Fig. 5h**) confirmed a reduction in nuclear neoplastic JUNB intensity upon TNF-α treatment (**Fig. 5i**). In addition, the expression of the CLA marker ECAD as well as nuclear GATA6 were significantly reduced (**Fig. 5j-l**). Together, these data indicate that TNF-α has direct implications in remodeling of the immune microenvironment and the plasticity of the neoplastic cells promoting PDAC aggressiveness.

**Figure 5.**
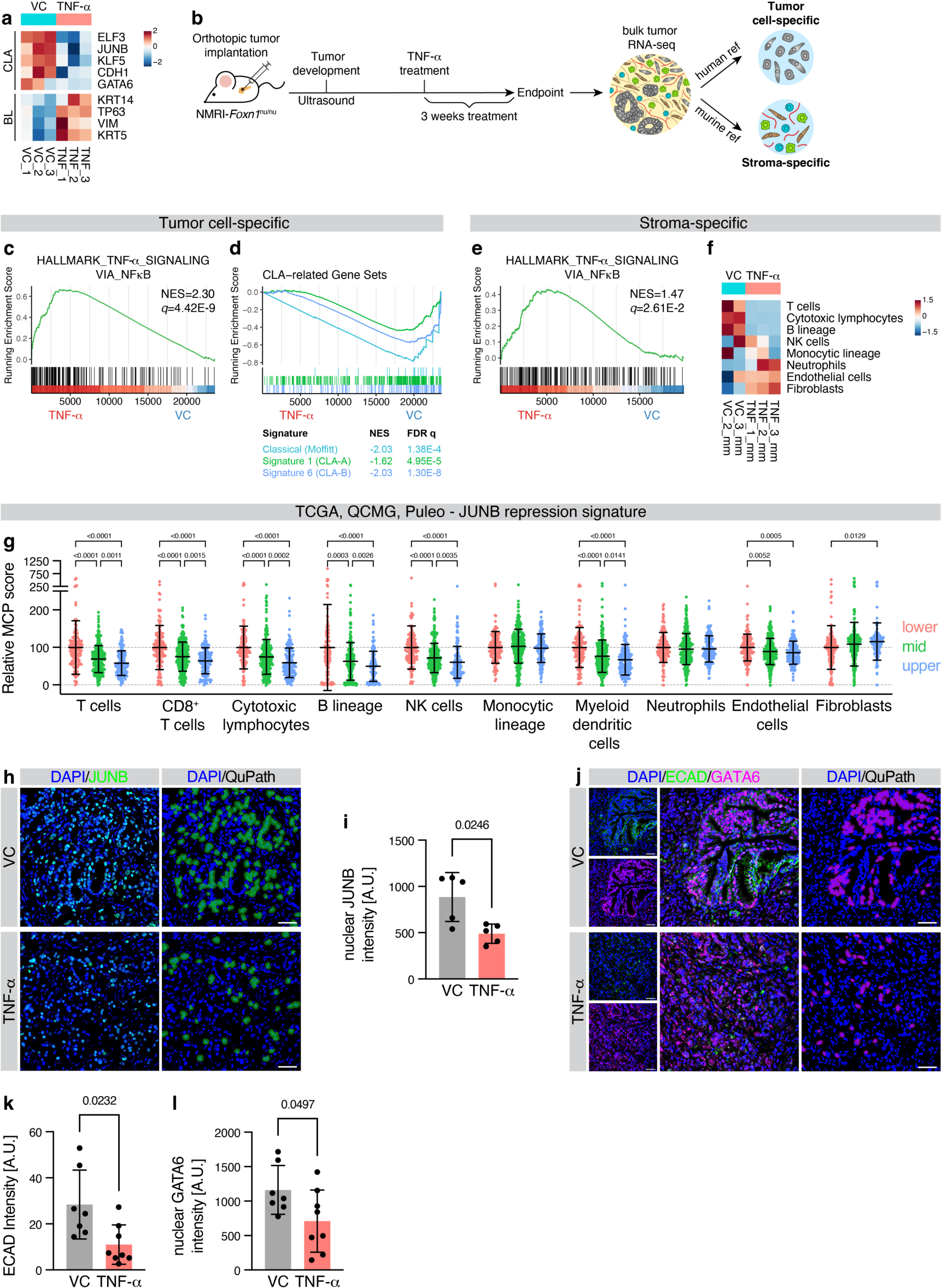
TNF-α disrupts CLA subtype identity and anti-tumor immunity. **a**, Heatmap of CLA and BL PDAC identity, in previously published^21^ RNA-seq data of CAPAN1 cells treated with TNF-α or vehicle control (VC) for 18 h. Cell color indicates z score. n=3. **b**-**f**, Virtually microdissected RNA-seq data of orthotopically transplanted CAPAN1 tumors treated with TNF-α or VC for three weeks. n=3 tumors; one stroma-specific transcriptome was excluded from the analysis. **b**, Flash-frozen tumors from orthotopically transplanted CAPAN1 tumors in NMRI-*Foxn1*^nu/nu^ mice treated with TNF-α or VC were bulk RNA-sequenced and subsequently aligned to human and murine reference genomes, to generate tumor and stromal cell-specific transcriptomes (Methods). **c**,**d**, Tumor cell-specific transcriptome. Gene set enrichment analysis (GSEA) for Hallmark signatures of the Molecular signature database (MSigDB) (**c**) and PDAC subtype signatures (**d**), for TNF-α versus VC. Normalized enrichment score (NES) and FDR *q*-value are indicated. **e**, As in **c**, for stroma-specific transcriptome. **f**, MCPcounter analysis in stroma-specific transcriptome. Cell color indicates z score. **g**, Relative MCPcounter scores for the indicated lineages in 582 patients of the TCGA, QCMG and Puleo cohort, separated into quartiles based on the JUNB repression signature score (as in **Figure 2j-m**). MCPcounter scores were min-max normalized and standardized to the mean of the lower JUNB repression signature score group for merging of the different cohorts. Mean ± s.d. shown. **h**, IF for JUNB in orthotopically transplanted CAPAN1 tumors treated with TNF-α or VC, with overlayed cell detection for nuclear JUNB^+^ cells by QuPath. Scale bar 50 μm. **i**, Quantification of **h**, for per-animal average nuclear JUNB intensity with mean ± s.d. shown. n=5. **j**, As in **h**, for ECAD and GATA6 staining and overlayed cell detection for nuclear GATA6^+^ cells. **k**, Quantification of **j** for per-animal average ECAD intensity per FOV with mean ± s.d. shown. **k**, As in **i**, for nuclear GATA6 intensity of **j**. **j**,**k**, VC, n=7; TNF-α, n=8. **g**,**i**,**k**,**l**, Student’s t-test with Welch’s correction.

### TNF-α promotes reactive spatial TiME heterogeneity in PDAC patients

Exogenous TNF-α treatment affected the TiME as well as neoplastic cell plasticity in experimental models (**Fig. 5**). Next, we set out to correlate TNF-α expression with spatial TME functions such as recruitment of CD68^+^ macrophages or CD3^+^, CD4^+^, CD8^+^ T cell infiltrations at histological levels in PDAC patients. We analyzed TNF-α expression and its effects in 105 PDAC patients by IHC. Overall, TNF-α levels were highly heterogeneous, with 46.9% of tumors displaying strong spatial variation in its expression (**Fig. 6a**). A major source of TNF-α in the TiME are macrophages^21^; indeed, TNF-α^high^ samples exhibited higher CD68 scores, both globally as well as in the individual samples (**Fig. 6b-d**). Importantly, TNF-α-dependent regional remodeling of the TiME seen in mice was recapitulated in patients, as the lymphocyte populations, particularly CD8^+^ T cells, were significantly reduced in TNF-α^high/int^ compared to TNF-α^low^ patients (**Fig. 6e, Extended Data Fig. 4a-g**). Recently, we have shown that the heterogeneous PDAC ecosystem self-organizes into ‘deserted’ and ‘reactive’ sub-tumor microenvironments (subTMEs), which leads to intratumoral zonation with co-existing immune-cold and immune-hot regions in human PDAC^35^. In accordance with these diverse tumor ecosystems, regional TNF-α expression was strongly increased within the immune-rich ’reactive’ subTME regions (**Fig. 6f-h**), which supports a BL aggressive phenotypic state in PDAC patients^35^. Together, these results indicate that high TNF-α levels are involved in an immunosuppressive TME that supports a BL state in PDAC.

**Figure 6.**
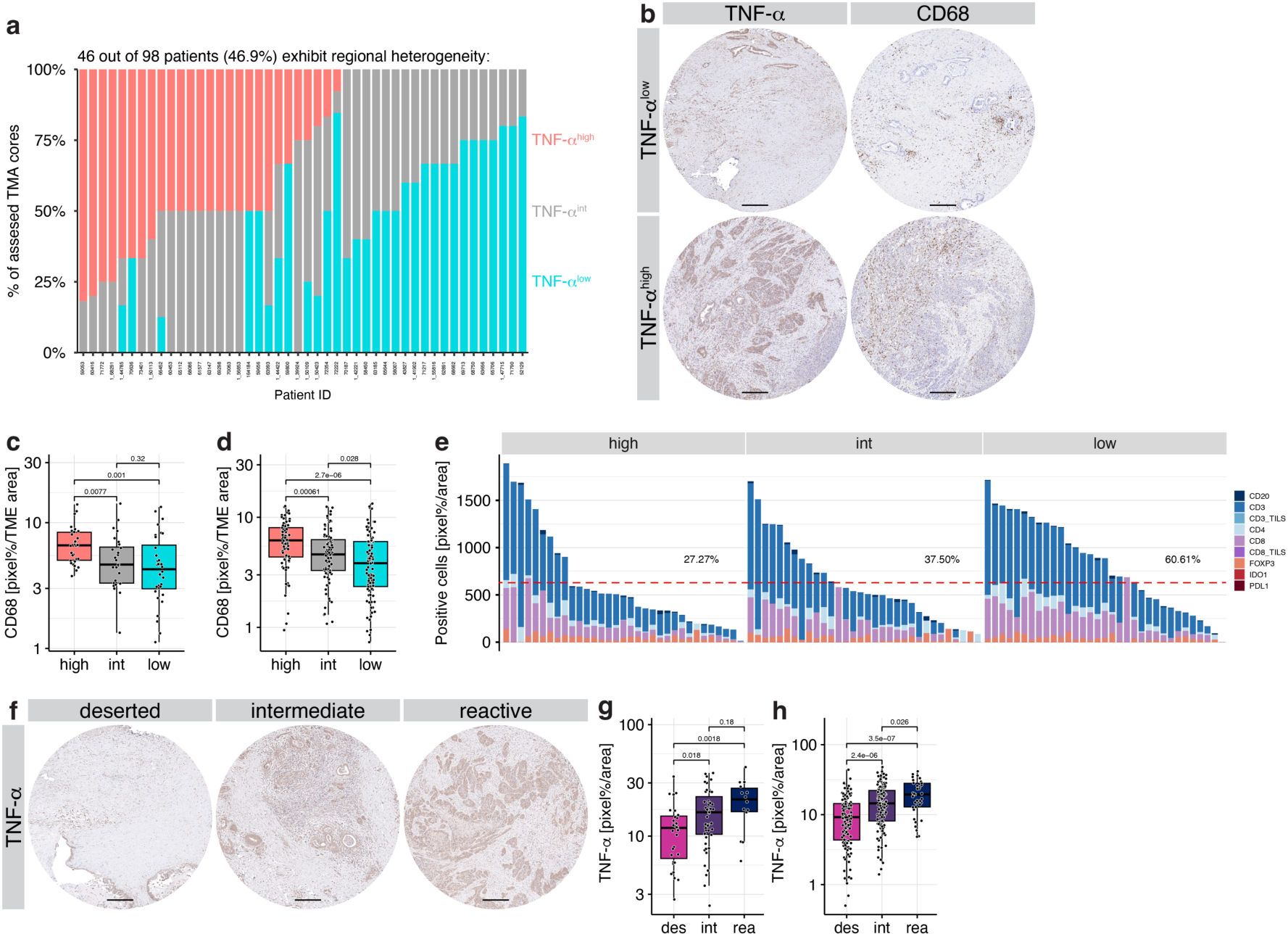
Spatial TNF-α expression promotes macrophage infiltration and T cell exclusion. **a**-**j**, IHC analysis in 105 PDAC patients or TNF-α expression. **a**, Spatial heterogeneity of TNF-α expression within different TMA cores of each patient. **b**, IHC for TNF-α and CD68 in cores classified as TNF-α^low^ and TNF-α^high^. Scale bar 200 μm. **c**,**d**, Quantification of **b**, in TNF-α^low^, TNF-α^intermediate^ (TNF-α^int^), and TNF-α^high^ expression per patient (**c**) and per TMA core across all patients (**d**). **e**, Lymphoid compartment distribution in TNF-α^low/int/high^ patients. Line and percentages denote patients above a third of the maximum value. **f**, Representative IHC staining of TMA cores for TNF-α in deserted, intermediate, and reactive subTMEs. Scale bar 200 μm. **g**,**h**, Quantification for TNF-α per patient (**g**) or TMA core (**h**) classified as deserted, intermediate, and reactive.

### Targeting TNF-α during chemotherapy leads to favorable TiME reorganization

Finally, we tested whether targeting TNF-α could shift tumors towards a favorable clinical state, given the central role of TNF-α in shaping PDAC subtype co-existence by influencing the JUNB-cJUN dichotomy. Anti-TNF-α monotherapy is not effective in aggressive PDAC^21^. Similarly, gemcitabine (GEM) chemotherapy alone or in combination with paclitaxel is essentially ineffective in *Kras*^G12D^;*p53*^R172H^;*Pdx1-Cre* (KPC)-derived murine PDAC models^27,38^. Thus, we tested whether combination of gemcitabine (GEM) with TNF-α inhibition may enhance treatment response. We utilized a highly aggressive KPC-derived orthotopic model and treated the animals with a combination of GEM plus anti-TNF-α monoclonal antibody therapy (**Fig. 7a**). This significantly prolonged overall survival from 19 to 32 days (**Fig. 7b**). While tumors maintained comparable gross histology (**Fig. 7c**), analysis of CD45, CD68, and TNF-α by IF showed a significant reduction in the number of CD45^+^/CD68^+^ macrophages, as well as reduction in CD45^+^/TNF-α^+^ cells (**Fig. 7d-f**). Notably, this also resulted in a significant increase in CD3^+^ as well as cytotoxic CD8^+^ T cells in the TME (**Fig. 7g-i**). Thus, TNF-α-dependent macrophage recruitment appeared to be halted, leading to a less immunosuppressive TiME in GEM/anti-TNF-α-treated PDAC tumors, which highlights the important role of TNF-α in shaping the immune landscape to aid tumor growth and survival. The central findings are summarized in **Fig. 7j**.

**Figure 7.**
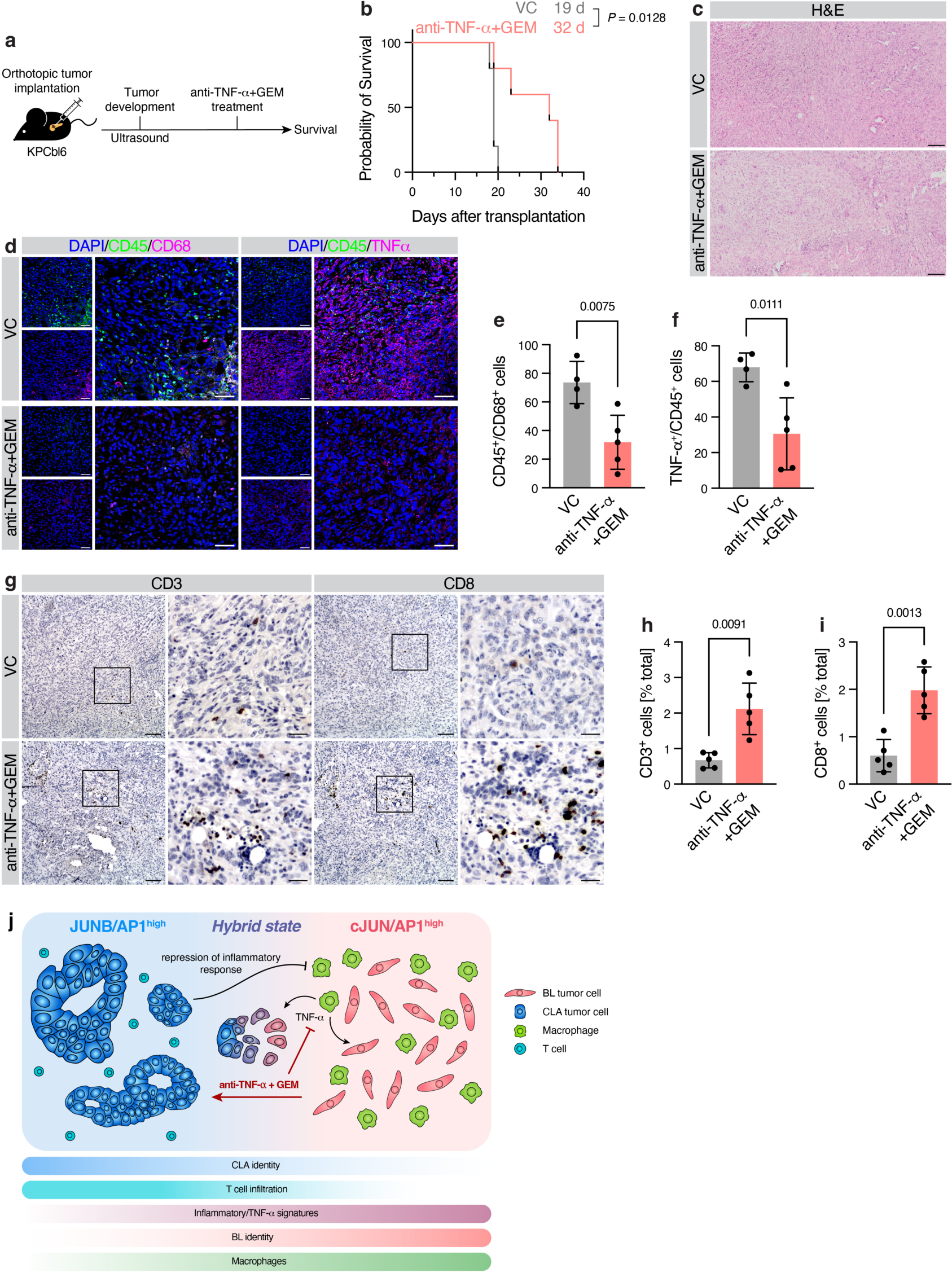
Targeting of TNF-α during chemotherapy restores anti-tumor immunity and prolongs survival. **a**, KPC cells were orthotopically implanted into syngeneic C57BL6/J mice and treated with an anti-TNF-α antibody in combination with gemcitabine (GEM) chemotherapy, or vehicle control (VC). **b**, Kaplan-Meier survival analysis of **a**. Median survival indicated. Log-rank test. **c**, H&E staining of anti-TNF-α+GEM and VC tumors. Scale bar 100 μm. **d**, IF for CD45 with CD68, and CD45 with TNF-α, in anti-TNF-α+GEM and VC tumors. Scale bar 50 μm. **e**,**f**, Quantification of **d** for CD45/CD68 (**e**) and TNF-α/CD45 (**f**) double-positive cells. Per-animal average counts per FOV with mean ± s.d. shown. **g**, IHC for CD3 and CD8 in orthotopically transplanted anti-TNF-α+GEM and VC tumors. Scale bar: overview, 100 μm; insert, 30 μm. **h**,**i**, Quantification of **g** for CD3^+^ (**h**) and CD8^+^ (**i**) cells. Per-animal average percentage of positive cells with mean ± s.d. shown. **e**,**f**,**h**,**i**, Student’s t-test with Welch’s correction. **i**, Model of AP1 dichotomy in PDAC subtype identity and immune recruitment.

## Discussion

PDAC is a highly heterogeneous disease, not only due to the intratumoral co-existence of neoplastic subtypes, but also in terms of its complex, overabundant TME. This subtype co-existence is increased during disease progression and negatively impacts both the predictive and prognostic utility of the transcriptomic subtypes. However, specific regional drivers of subtype identity and their potential relationship with the heterogeneous TME are currently unknown.

Here, we investigated the role of neoplastic AP1-mediated epigenetic and transcriptional programs in shaping the local inflammatory TiME, which in turn is critical for intratumoral subtype plasticity and PDAC aggressiveness^10–15,20,36,37^. We report that AP1 transcription factors (JUNB/AP1 vs. cJUN/AP1) hold a dichotomous role in maintaining both the plasticity and stability of CLA and BL neoplastic cells via intrinsic epigenetic and transcriptional regulation of lineage gene expression as well as extrinsic inflammatory processes. Integrated bioimaging, epithelial-specific transcriptome, and LCM-multiOME analyses of PDAC patients showed that high JUNB expression is associated with GATA6^+^ CLA identity. Mechanistically, neoplastic JUNB positively controls the regulation of CLA-specific lineage factors (HNF1B and GATA6), while epigenetically repressing BL-specific inflammatory immune regulators (cJUN), which is critical for the maintenance of the CLA neoplastic identity. Intriguingly, this JUNB-mediated CLA subtype differentiation is not stable, but highly plastic in response to inflammatory cues (i.e., TNF-α) or chemo-radiation therapy, as characterized by loss of JUNB^+^/GATA6^+^ cells in preclinical models and in PDAC patient specimens. These findings clearly indicate that loss of JUNB/AP1-dependent gene regulation leads to destabilization of CLA neoplastic identity, induces a CD68^+^/TNF-α^+^ macrophage-driven inflammatory response in the TiME and, thereby, promotes a BL invasive state via complementary intrinsic and extrinsic mechanisms. Specifically, the TNF-α-mediated inflammatory response associated with low JUNB expression destabilizes CLA neoplastic identity by promoting a CLA-to-BL transition via epigenetic transcriptional reprogramming in PDAC. Altogether, these results unveil a key reciprocal interdependence between neoplastic (intrinsic) and local microenvironmental (extrinsic) factors that influence subtype plasticity/instability and thereby promote PDAC heterogeneity and aggressiveness.

Emerging evidence shows that cJUN/AP1 transcription factors play an important role in tumor inflammation, chemotherapy response and tumor recurrence in PDAC patients^21,32^. Complementarily, we here show that JUNB/AP1 acts as a counterpart to promote a favorable CLA phenotypic identity in PDAC. Of note, JUNB/AP1-mediated transcriptional programs can also confer tumor-promoting functions in other cancer types^39–41^; JUN/AP1 TFs are highly context dependent and may co-operate for target gene transcription^41–43^ or oppose one another^44^. Our data indicates that in PDAC, JUN/AP TFs exert antagonistic roles, with JUNB directly repressing cJUN and cJUN-regulated cytokine secretion, thereby inhibiting recruitment of CD68^+^/TNF-α^+^ macrophages in the TME. Thus, JUNB not only maintains a CLA subtype lineage with high expression of epithelial differentiation factors such as GATA6, but also directly represses drivers of disease aggressiveness (e.g., MYC) as well as inflammatory pathways associated with poorer clinical outcome in PDAC patients. In this context, our model of CLA-derived orthotopic tumors with cJUN-OE provides a key experimental benefit in that it recapitulates the intratumoral subtype coexistence seen in PDAC patients. Specifically, this model revealed significant infiltration of CD68^+^/TNF-α^+^ macrophages in the spatial tumor neighborhood of cJUN^high^ but not cJUN^low^ neoplastic cells, indicating that cJUN exploits regional macrophages to attenuate JUNB expression and thereby suppress CLA neoplastic identity. These findings were crucial in revealing the mechanisms maintaining AP1 dichotomy: direct (JUNB repressing cJUN) and indirect (cJUN repressing JUNB via local macrophages), which regulate the divergent expression and functions of the AP1 TFs in a reciprocal manner with the spatial TiME.

Altogether, this string of insights provides a potential mechanistic foundation for several recent studies that showed a high degree of heterogeneity in the neoplastic and stromal immune compartments in human PDAC, including hybrid/intermediate/co-expressor CLA/BL subtype states that exist in naive and therapy-treated PDAC tumors^10–16,20,36,37^. We propose that extrinsic regional TNF-α plays an essential role in destabilizing CLA neoplastic cell identity by promoting BL cJUN/AP1-mediated transcriptional programs. Compartment-specific transcriptomic profiling of TNF-α-treated tumors demonstrated that TNF-α strongly affects both neoplastic-specific as well as immunosuppressive stromal gene expression. In the neoplastic compartment, TNF-α treatment induced loss of CLA identity, whereas in the TME it resulted in depletion of T and B cell signatures. In line, in a large cohort of PDAC patient tumors, we observed a significant reduction of CD3^+^, CD4^+^, and CD8^+^ T cell infiltrations particularly in TNF-α^high^ tumors, whereas CD68^+^ macrophages were strongly elevated, underlining how recruitment of inflammatory CD68^+^ macrophages to BL TNF-α^high^ regions simultaneously leads to an immunosuppressive TME. Furthermore, we also found strongly enhanced TNF-α expression in reactive subTME regions^35^, strengthening TNF-α as a major link between the TME and the BL inflammatory subtype, as reactive subTMEs provide the organizational framework for an inflamed, poorly differentiated and aggressive tissue state with CK5^high^/GATA6^low^ neoplastic cells. To thus test the potential utility of TNF-α in new avenues of therapeutic interference, we treated a highly aggressive KPC tumor model with an anti-TNF-α monoclonal antibody plus GEM. Combined anti-TNF-α and chemotherapy substantially improved the survival, reduced the infiltration of CD45^+^/CD68^+^/TNF-α^+^ macrophages, and induced recruitment of CD3^+^ and CD8^+^ T cells in the TME. This identifies novel TNF-α-specific functions that appear to determine an immunosuppressive regional TiME and PDAC aggressiveness. In accordance, the recent evaluation of the PRINCE trial indicates that while higher CD4^+^/8^+^ T cells are associated with better response to immune checkpoint inhibition with chemotherapy, elevated TNF-α signaling negatively impacts therapy response in metastatic PDAC patients^45^, further corroborating a major role of TNF-α in immunosuppression.

In sum, our comprehensive analysis of the dichotomous role of the AP1 TFs in PDAC subtype heterogeneity has shed light onto the mechanism of JUNB/HDAC-dependent suppression of BL-associated inflammatory responses. JUNB signaling can be regionally repressed through macrophage-secreted TNF-α, destabilizing favorable CLA subtype identity and inducing T cell exclusion, which highlights the important role of the single cytokine TNF-α in shaping an immunosuppressive TiME and tumor aggressiveness. Thus, therapeutically shifting the balance from T cell^low^/macrophage^high^/BL towards T cell^high^/macrophage^low^/CLA state through inhibition of TNF-α with GEM chemotherapy may provide a valuable strategy to enhance anti-tumor immunity and treatment response in PDAC.

## Methods

### Preclinical animal experiments

All animal experiments were conducted at the UMG following Central Animal-experimental authority guidelines (permission no. 15/2057, 14/1634, 18/2953). The ethics committee of the UMG also approved the generation of the PDX mouse model (permission no. 70112108). Eight to ten-week-old NMRI-*Foxn1*^nu/nu^ or C57BL/6J mice were used for orthotopic transplantation as previously described^21^. 1 × 10^6^ cells (CAPAN1, CAPAN2, CAPAN2-EV, or CAPAN2-cJUN) were transplanted orthotopically into NMRI-*Foxn1*^nu/nu^ mice, or 3.5 × 10^4^ KPC cells into syngeneic C57BL/6J mice, each in 15 μL culture medium under isoflurane anesthesia. Tumor growth was monitored by weekly ultrasound. The TNF-α treatment model in orthotopic CAPAN1 tumor model has been described previously^21^. For the anti-TNF-α plus gemcitabine treatment model, C57BL/6J mice bearing orthotopic KPC tumors were treated with 10 μg g^−1^ anti-TNF-α antibody (BioLegend) in combination with 100 mg/kg gemcitabine (Sigma-Aldrich) three times a week for three weeks. Preclinical studies were terminated when mice displayed exclusion criteria, e.g. body weight loss >20%, tumor diameter of >15 mm, or overall poor clinical presentation. Tissues were fixed in formalin and embedded in paraffin for histological analysis.

### Patient data analysis and study approval

For correlation of gene expression in patient tumors, publicly available datasets^4,33,34^ as well as previously described compartment-sorted patient transcriptomes^22^ were used. For the compartment sorted transcriptomes, in brief, tumor tissue of untreated patient with partial pancreatoduodenectomy at the Department of General, Visceral and Transplantation Surgery, University of Heidelberg (HIPO-project approved by the ethical committee of the University of Heidelberg; case number S-206/2011 and EPZ-Biobank Ethic Vote no. 301/2001) were subjected to fluorescence-activated cell sorting with compartment-specific markers (for epithelial EPCAM^+^/CD45^−^/CD31^−^) and subsequently RNA-sequenced in the sorted fractions. TCGA^34^ and QCMG^4^ datasets were accessed via cBioPortal^46,47^, Puleo^33^ dataset was accessed via the ArrayExpress database with accession number E-MTAB-6134. The expression values were plotted as z scores (TCGA), RMA-normalized probe intensities (of the highest-expressing probe for one gene; Puleo), log(RSEM) (QCMG), or log2(TPM+1) (compartment-sorted patients). Survival data was available for 288 of the total 309 patients of the Puleo cohort.

Transcriptome and proteome data of LCM-enriched PDAC tumor epithelia and stroma has been described previously^35^. In brief, cryosections of 32 PDAC patient tumors were microdissected for epithelia and stroma, and separately approx. 50000 to 100000 cells prepared for RNA-seq and shotgun proteomics each. R v4.2.0 was utilized for the analysis. For correlation analysis, Spearman’s *R* and *P* value, as well as linear regression with 95% CI are indicated in the figures, plotted using the ggscatter function of the ggpubr package v0.5.0. GSEA analysis of compartment-sorted patient data was performed as described for RNA-seq data, using a gene list sorted by Spearman’s *R* for correlation of all genes to JUNB as input. JUNB repression signature scores were calculated using the GSVA package^48^ v1.44.5 for expression data of each cohort individually. MCPcounter scores were determined with the MCPcounter package as above. For merging of the MCPcounter scores of the different cohorts, scores were min-max normalized and standardized to the mean of the lower quartile of the JUNB repression signature scores. Patient survival analysis was performed using the survival v3.5-5 and ggsurvfit v0.3.0 packages. Expression heatmaps and hierarchical clustering were created by the pheatmap package v1.0.12.

### Human PDAC tissue IF and tumor microarray IHC analysis

PDAC resection tissue utilized for IF staining was derived from the Molecular Pancreatic Cancer Program (MolPAC) of the University Medical Center Göttingen as part of the study “Klinische und molekulare Evaluation von Patienten mit Pankreasraumforderungen im Rahmen des Pankreasprogramms der UMG (MolPAC)” (ethics approval 11/5/17). IF staining was performed as described above.

The PDAC tissue microarray (TMA) cohort has been described previously with details on analyses and overall staining detailed in the original publication^35^. In addition to the described staining, IHC was performed for JUNB (Cell Signaling #3753) and TNF-α (abcam ab1793) manually according to standard laboratory procedures using antigen retrieval by Tris-EDTA buffer solution (pH 9.0). Quantification was performed using QuPath measuring positive pixel percentages. Epithelial tumor-specific areas were annotated manually.

### Cell culture

Established human PDAC cell lines CAPAN1 and CAPAN2 were purchased from ATCC (Manassas, VA) and authenticated (CAPAN1, RRID:CVCL_0237), (CAPAN2, RRID:CVCL_0026) by the German Collection of Microorganisms and cell culture GmbH (DSMZ). PDAC cell lines were cultured in RPMI 1640 (Thermo Fisher Scientific) with 10% (v/v) fetal calf serum (FCS; Th. Geyer). Murine cells derived from the *Kras*^G12D^;*p53*^R172H^;*Cre* mouse model (“KPC cells”) were maintained in DMEM with 10% FCS and 1% non-essential amino acids. Patient-derived primary cell line GCDX62 was maintained in a 3:1 mixture of Keratinocyte-SFM (KSF; Thermo Fisher Scientific; supplemented with 2% (v/v) FCS, 1% (v/v) Penicillin-streptomycin, bovine pituitary extract (BPE), and human epidermal growth factor) and RPMI 1640 containing 10% (v/v) FCS. cJUN overexpression (cJUN-OE) and empty vector (EV) control cell lines of CAPAN2 and GCDX62 were generated as previously described^21^ and maintained in their normal growth medium supplemented with 1 μg/mL puromycin.

### siRNA transfection

For transient knockdown experiments, 5 × 10^5^ cells were seeded in 6-well plates and immediately transfected with a mixture of 10 μL Lipofectamine2000 (Thermo Fisher Scientific), 6 μL of 20 μM target-specific siRNA (or non-targeting siRNA as control), and 200 μL Opti-MEM (Thermo Fisher Scientific) after 15 min incubation of the transfection mixture at room temperature (RT). Cell culture medium was changed after 24 h, and protein or RNA extracted 48-72 h after transfection.

### Immunoblotting

Cells were washed with phosphate-buffered saline (PBS) and lysed in whole cell lysis (WCL) buffer supplemented with cOmplete protease inhibitor cocktail (Roche Diagnostics), 100 μM sodium orthovanadate (NaO; Sigma-Aldrich) and 100 μM phenylmethylsulfonyl fluoride (PMSF; Sigma-Aldrich) for 30 min on ice. Cell lysates were centrifuged at 17000 × g for 20 min at 4°C, and supernatants collected. Proteins were diluted to 1 μg/μL in WCL and Laemmli buffer and boiled for 8 min at 95°C. Denatured samples were resolved by SDS-PAGE on 10 or 15% gels and transferred onto nitrocellulose membranes. After blocking in 5% (w/v) milk powder in TRIS-buffered saline with 0.1% (v/v) Tween 20, primary antibodies were incubated overnight at 4°C. Secondary HRP-linked antibodies were subsequently incubated at RT for 1 h. Next, membranes were developed with ECL solution using a ChemoStar imager (Intas Science Imaging Instruments). Antibodies are listed in **Supplementary Table 1**.

### Co-immunoprecipitation

Cells were seeded in 10 cm dishes and harvested by scraping in 1.5 mL ice-cold PBS. The cell suspension was centrifuged at 500 × g for 5 min at 4°C. The resulting pellet was resuspended in lysis buffer containing Triton X-100. After the cells were completely lysed, the lysates were centrifuged at 17000 × g for 20 min at 4°C and the supernatant transferred into new tubes. 500 μg of protein were added to washed agarose protein A beads (Merck Millipore) and incubated on a rotating wheel for 1 h at 4°C. Afterwards, beads were removed by centrifugation and precleared lysates were incubated with the target antibodies overnight at 4°C. The following day, 50 μL of washed agarose A beads were added to the lysates and incubated on a rotating wheel for 2 h at 4°C. The beads/antibody/target complexes were washed twice with WCL buffer and twice with PBS with cOmplete protease inhibitor cocktail. Finally, the complexes were resuspended in 65 μL 2x Laemmli buffer and incubated for 8 min at 95°C. These samples were processed for immunoblotting as described above. To avoid the heavy chain signal of the pulldown antibody in case pulldown and primary immunoblot antibody were of the same species, an anti-light-chain antibody raised in another species was used prior to using the secondary antibody. Antibodies are listed in **Supplementary Table 1**.

### RNA isolation and quantitative real-time PCR

Total RNA was extracted using TRIzol reagent (Invitrogen) according to the manufacturer’s protocol. Briefly, cells were washed with ice-cold PBS and collected in 800 μL TRIzol reagent, followed by addition of 200 μL chloroform after a short incubation. The solution was vortexed for 5 seconds to mix it thoroughly. After incubation at RT for 5 min, samples were centrifuged at 17000 × g for 15 min at 4°C. Next, the upper aqueous phase was transferred into a new 1.5 mL tube. Subsequently, 500 μL of isopropanol was added and mixed. After centrifugation at 17000 × g for 30 min at 4°C, the resulting pellet was washed twice with 75% ethanol. Finally, the dried pellet was dissolved in 30 μL of nuclease-free water. 1 μg RNA was used for cDNA synthesis using the iScript cDNA Synthesis Kit (BioRad) according to the manufacturer’s instructions. 5 μL of SYBR green (BioRad) and 0.25 μL of forward and reverse primers each were mixed with 1 μL of cDNA. Quantitative real-time PCR (qRT-PCR) was performed using the StepOnePlus Real-Time System (Applied Biosystems). Relative quantification values were calculated with the associated StepOnePlus software, using *XS13* as reference control gene. Primer sequences are listed in **Supplementary Table 2**.

### Chromatin immunoprecipitation

For chromatin immunoprecipitation followed by quantitative real-time PCR (ChIP-qPCR), cells were seeded in 10 cm culture dishes and fixed in 1% PFA (Thermo Fisher Scientific) in PBS for 20 min at RT. After quenching with 1.25 M glycine, cells were washed in PBS and scraped in 1.5 mL cold Nelson buffer with cOmplete protease inhibitor cocktail, 100 μM NaO and PMSF, and 10 mM sodium fluoride . Following centrifugation at 12000 × g for 2 min at 4°C and washing with Nelson buffer with inhibitors, nuclear pellets were snap-frozen in liquid nitrogen. Nuclear pellets were lyzed in Gomes lysis buffer with cOmplete protease inhibitor cocktail, 100 μM NaO and PMSF, and 10 mM sodium fluoride with 0.1% SDS (for JUNB, HDAC1) or 0.5% SDS (for H3K27ac). After lysis for 15 min at 4°C, cells were sonicated on a Bioruptor Pico (Diagenode) for 3-8 cycles with 30 seconds ON/OFF. Sonication efficiency was validated by agarose gel electrophoresis. Thereafter, lysates were pre-cleared with 15 μL washed protein A/G magnetic beads (Thermo Fisher Scientific) and then incubated over night with the primary pulldown antibodies or isotype control (**Supplementary Table 1**). For pulldown of antibody-antigen complexes, 30 μL of washed beads are added to lysates and incubated for 2 h at 4°C. Finally, lysates are washed with Gomes lysis buffer, Gomes wash buffer and TE buffer. Corresponding input and pulldown samples are then RNase A and proteinase K digested. DNA was isolated by phenol-chloroform-isoamyl alcohol method. RT-qPCR was performed as described above, increasing total PCR cycles to 55. Primer sequences are listed in **Supplementary Table 2**.

### RNA-seq and ChIP-seq analysis

For RNA-seq, library preparation and sequencing for CAPAN1 cells subjected to JUNB silencing (siJUNB) or non-targeting control (siCtrl) were performed as described previously^21^. For comparability, previously published RNA-seq data of TNF-α-treated CAPAN1 cells was analyzed analogously using the following pipeline.

Raw reads were quality-checked using FastQC, aligned to hg38 reference genome using STAR^49^ v2.7.3a and counted per gene using htseq-count^50^ v0.11.3. Downstream analysis was conducted in R v4.2.0 and Bioconductor^51^ packages v3.15. Variance stabilization was performed using RUVSeq^52^ v1.30.0 function RUVs. Differential gene expression was performed using DESeq2^53^ v1.36.0. Gene set enrichment analysis^54^ (GSEA) was conducted with clusterProfiler^55^ v4.4.4, using signatures of the Molecular Signature Database^56^, as well as custom signatures based on published PDAC subtype classifications^4,5,12,57^.

For tissue RNA of TNF-α or VC-treated tumors of orthotopically implanted CAPAN1 cells, RNA was purified using Direct-ZOL RNA Mini-prep (Zymo Research). Library preparation and sequencing (paired-end, 150 bp read length) was performed by Novogene. To allow deconvolution of human (tumor cell) and murine (host stroma) compartments, reads were aligned using STAR v2.7.3a to human reference hg38, as well as murine reference mm39. Subsequently, the XenofilteR^58^ package v1.6 in R v4.1.0 was used to remove either murine reads from the human alignment or *vice versa*. After filtering, samples were processed as above.

To estimate the relative abundance of stromal cells, murine reads were processed with the MCPcounter^59^ package v1.2.0 in R v4.2.0.

JUNB and cJUN ChIP-seq, as well as ATAC-seq, were conducted previously^21^, accessible at GSE179781. Further, previously published RNA-seq data for GCDX62-cJUN-OE and GCDX62-EV was utilized, which is accessible at GSE173121. For meta-pathway analysis of JUNB-bound and accessible regions, genes were annotated to JUNB-bound and ATAC-seq peak regions as described previously and analyzed using Metascape^60^ (https://metascape.org/). For integration of RNA- and ChIP-seq data, consensus ChIP-seq peaks were annotated to genes using the R package rGREAT^61^ v3.0.0, and their fold change in the corresponding RNA-seq extracted. Gene ontology (GO) analysis was then performed for the significantly up- or downregulated genes of this subset using the clusterProfiler package as above. The full list of JUNB-bound genes that are significantly upregulated upon siJUNB, termed the JUNB repression signature, is available in **Supplementary Table 3**.

### Reporter assay

For reporter assays, the Dual-Luciferase Reporter Assay System (Promega) was used. 5 × 10^4^ cells were seeded into 24-well plates. The following day, cells were transfected with 200 ng of pJC6-GL3 (#11979, Addgene; cJUN), which contains the cJUN promoter linked to the firefly luciferase gene. Additionally, cells were co-transfected with different concentrations of JUNB-overexpressing plasmids (Paul Dobner, University of Massachusetts), 15 ng of pRL-null *Renilla* luciferase control reporter (#E2271, Promega; for background normalization), and compensated with empty vector control (pCMV2c, David Russell, UTSW) to ensure equal total amounts of transfected plasmids. Reporter, overexpression, *Renilla* luciferase and empty vector plasmid mixture was added to Lipofectamine 2000 (Thermo Fisher Scientific) and 50 μL of Opti-MEM (Thermo Fisher Scientific) and incubated at RT for 10 min. Thereafter, 50 μL of the transfection mixture was added to each well for 24 h. Before the measurement, cells were lysed with 1x Passive Lysis Buffer (Promega) for 10 min on a shaker at RT. Subsequently, 30 μL of the cell lysate was transferred into a white 96-well plate and 30 μL of firefly luciferase substrate (Promega) was added. The luminescence was measured using a LUmo microplate reader (Autobio Diagnostics). Next, 30 μL of the Stop & Glo reagent (Promega) was added to the previous solution and luminescence measured again for the *Renilla* signal. Firefly luciferase luminescence signal was finally divided by the *Renilla* luciferase luminescence and normalized to the control sample.

### Transwell invasion assay

For invasion assays of silenced cells, siRNA transfection was performed 24 h before as described above. 8 μm porous cell culture inserts (Falcon) were coated in type I collagen (Enzo; diluted 1:62.5 in 0.1 M HCl) for 2 h. Then, silenced or control cells were seeded in 50 μL Matrigel solution (Corning) and solidified for 30 min before adding normal culture medium to the inserts and wells. After 48 h incubation, medium was aspirated, Matrigel removed, and membranes fixed in 4% PFA. After washing with PBS, membranes were stained with DAPI for 1 min and finally mounted on glass slides in Immu-Mount (Thermo Fisher Scientific). Invaded cells were imaged using a DMi8 fluorescence microscope (Leica) and quantified manually using ImageJ Fiji^62^. For each biological replicate, two independent inserts were evaluated.

### Hematoxylin and eosin staining

Hematoxylin and eosin (H&E) staining was performed as previously described^21^. Formalin-fixed paraffin-embedded tissues were cut into 4 μm thin sections. Tissue sections were incubated in xylene for 1 h and rehydrated in decreasing concentrations of ethanol and finally water. Then, sections were stained with hematoxylin for 8 min, followed by bluing for 7 min under running tap water. Subsequently, sections were briefly incubated in mild acetic acid solution and transferred into eosin/acetic acid solution for 3 min. Lastly, slides were dehydrated in an increasing ethanol series and mounted with Roti-Mount (Carl Roth).

### Immunofluorescence and immunohistochemical staining

Immunofluorescence (IF) staining was performed as previously described^21^. Briefly, slides were deparaffinized and rehydrated followed by antigen retrieval by boiling in citrate buffer (pH 6.0). Sections were blocked in 1% (w/v) bovine serum albumin (BSA; Sigma) in phosphate buffer (PB) containing 0.4% Triton X-100. After washing five times with PB, sections were incubated with primary antibodies at 4°C overnight. After six PB washes, fluorophore-coupled secondary antibodies were incubated at 4°C for 2 h. Subsequently, sections were washed in PB, stained with DAPI, and mounted in Immu-Mount (Thermo Fisher Scientific).

For immunohistochemistry (IHC) staining, the VECTASTAIN ABC-HRP Kit and ImmPACT DAB Substrate Kit (Vector Laboratories) was used. Deparaffinization and antigen retrieval was performed as above. Then, slides were fixed on a slide holder and incubated with 3% hydrogen peroxide solution for 10 mins prior to blocking in 10% BSA. Next, the slides were incubated with primary antibodies overnight at 4°C. The next day, slides were washed three times with PBS with 0.1% Tween20 (Sigma; PBST) and incubated with secondary antibodies for 1 h followed by AB complex incubation. Afterwards, slides were washed with PBST and stained with DAB solution and further placed in deionized water to stop the reaction. Slides were then stained with hematoxylin for 7 min and blued under running water. Lastly, the slides were dehydrated and fixed as described for H&E staining.

### Image acquisition and analysis

For bright-field applications (H&E, IHC), images were acquired using either an Olympus BX43 light microscope or the Olympus VS120 virtual slide microscope. IF staining was imaged using either an Olympus IX81 confocal fluorescence microscope, or the Olympus VS120 virtual slide microscope.

Quantification of IF and IHC images was performed either using ImageJ Fiji^62^ (by manually counting positive cells, measuring the fluorescence intensity of the entire field of view, or using semi-automatic macros as described previously^63^), or using QuPath^64^ v0.4.3. In QuPath, cell detection was performed using the built-in detection feature, obtaining intensity measurements per cell, per nucleus or per cytoplasm, which were used for positive cell quantification using single measurement classifications (or compound classifiers for double-positive cells). For IHC images, positive cell detection using optical density for nuclei detection and deconvoluted DAB intensity for positive cells was used.

For hotspot analysis of HA-cJUN IHC and JUNB IF, density maps of positive cells were created and above a threshold defined as high-density areas (“hotspots”), which were then transferred to the respective other slide image to derive positive cells within cJUN or JUNB hotspot regions. For distance analysis of hotspots to CD68^+^ cells, cell detection was performed as above, the hotspot regions transferred to the CD68 staining image, and the 2D distance to annotation tool of QuPath used.

### Statistical analysis

GraphPad Prism version 8.0.2 was used for statistical analysis. The comparison of independent groups was performed using an unpaired Student’s t-test with Welch’s correction. One-way analysis of variance (ANOVA) was used for more than three conditions of a single factor. Survival data was analyzed using the log-rank test. Results were considered significant with a *P* value below 0.05, as indicated in the figures. Spearman correlation coefficient with a two-tailed *P* value was used for the patient gene expression correlation data.

## Acknowledgements

This study was supported by the Deutsche Krebshilfe (70112999; 70115054; Max-Eder Program), the Fritz-Thyssen Stiftung (1842610) and the Wilhelm-Sander-Stiftung (2021.159.1) to SKS. This study was also supported by KFO 5002 grant of the DFG to EH, AP, VE and SKS. NK was funded by the DFG (413501650). The processing of human PDAC patient data was made possible by the Dietmar-Hopp Foundation and the BioRNSpitzen cluster ’Molecular- and Cell-based Medicine’; the German Ministry of Science and Education (BMBF) e:Med program for systems biology (PANC-STRAT consortium, grant no. 01ZX1305C and 01ZX1605C). The PancoBank collection and processing of human specimens were supported by Heidelberger Stiftung Chirurgie. We thank L. Schüürhuis, K. Reutlinger, S. Mercan, and W. Kopp for their technical assistance. We sincerely thank the EPZ-Biobank and Department of General and Visceral Surgery of the University Hospital Heidelberg, particularly N.A. Giese, T. Hackert, O. Strobel and M. Büchler for their collaboration involving fresh primary PDAC specimens.

## Author contributions

LK, MT, and SKS designed the overall study. SKS and LK wrote the manuscript with contributions of MT. BTG and UK edited the manuscript. MT performed preclinical studies. MT, LK, DG, and FP analyzed tissue of patients and preclinical models. MT, LU, and FP performed *in vitro* experiments. LK performed bioinformatic analysis of ChIP-, ATAC-, and RNA-seq data from *in vitro* and *in vivo* models, as well as analysis of patient expression data, with AP aiding ChIP- and ATAC-seq data analysis. EE performed RNA-seq of flow cytometry-sorted patient specimens, EE and LK analyzed the data, with EE and AT aiding in data interpretation. EH provided human PDX and patient specimens. SKS, EH, and VE critically analyzed murine and human histopathological examinations and data interpretation. NK, NC, KA, FY and BTG performed patient IHC analysis, BTG and RK helped with experimental design and data interpretation. SKS supervised the project and interpreted the data.

## Competing interests

The authors have no conflict of interest to declare.

## Data availability

For this study, the Molecular Signatures Database (https://www.gsea-msigdb. org/gsea/msigdb/) database was used. Previously published^21,23^ ChIP- and ATAC-seq data are available at Gene Expression Omnibus (GEO) under accession codes GSE173159 and GSE64560. RNA-seq data generated for this study has been deposited at GEO with the accession code GSE245941. FACS-sorted epithelial patient tumor RNA-seq data^22^ are available at EGA under accession code EGAS00001004660. Patient tumor LCM-enriched transcriptome and proteome data are available at EGA under accession code EGAS00001002543 and from UCSD’s MASSive database under accession code MSV000086812. Processed proteome data is detailed in the previous publication^35^.

All other data supporting the findings of this study are available from the corresponding author on reasonable request.

## Extended Data Figures

**Extended Data Figure 1.**
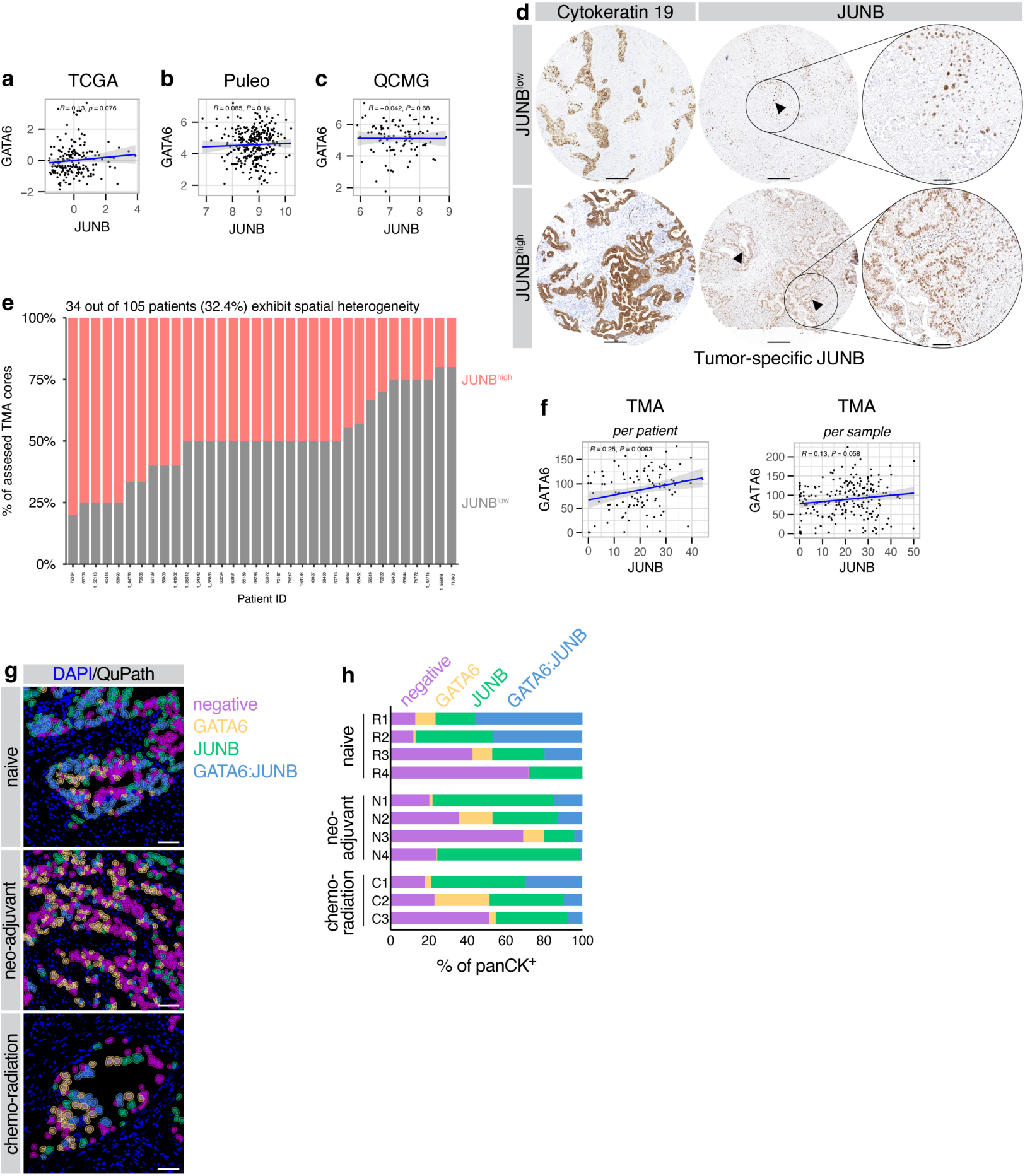
Association of cJUN with CLA PDAC markers in patients. **a**-**c**, Gene expression correlation of JUNB with GATA6 in the TCGA (**a**; n=177), Puleo (**b**; n=309), and QCMG (**c**; n=96) cohorts. Linear regression with 95% CI, as well as Spearman’s *R* and associated *P* value. **d**-**f**, IHC analysis in 105 PDAC patients for epithelial JUNB expression. **d**, IHC for cytokeratin 19 and JUNB in TMA cores classified as JUNB^low^ and JUNB^high^, indicating tumor-specific JUNB^+^ cells. Inserts show higher magnification of JUNB staining. Scale bar: overview 200 μm; insert 50 μm. **e**, Spatial heterogeneity of tumor-specific JUNB expression within different TMA cores of each patient. **f**, Correlation of JUNB and GATA6 IHC quantification per patient (left) or per TMA core across all patients (right), plotted as in **a**-**c**. **l**, Representative DAPI staining with QuPath overlay for cell detection of staining for JUNB, pan-cytokeratin (panCK), and GATA6, in tissue of treatment-naive, neo-adjuvant-treated, and chemo-radiation-treated PDAC patients (shown in Figure 1p). Classifications for cytoplasmic panCK^+^ cells that are additionally positive for nuclear GATA6 (yellow), nuclear JUNB (green), both nuclear GATA6 and JUNB (blue) or only panCK (“negative”; magenta), are indicated. Scale bar 50 μm. **h**, Quantification of **g** showing percentage of panCK^+^ cells. Each patient is shown individually as stacked bar graphs. Naive, n=4; neo-adjuvant, n=4; chemo-radiation, n=3.

**Extended Data Figure 2.**
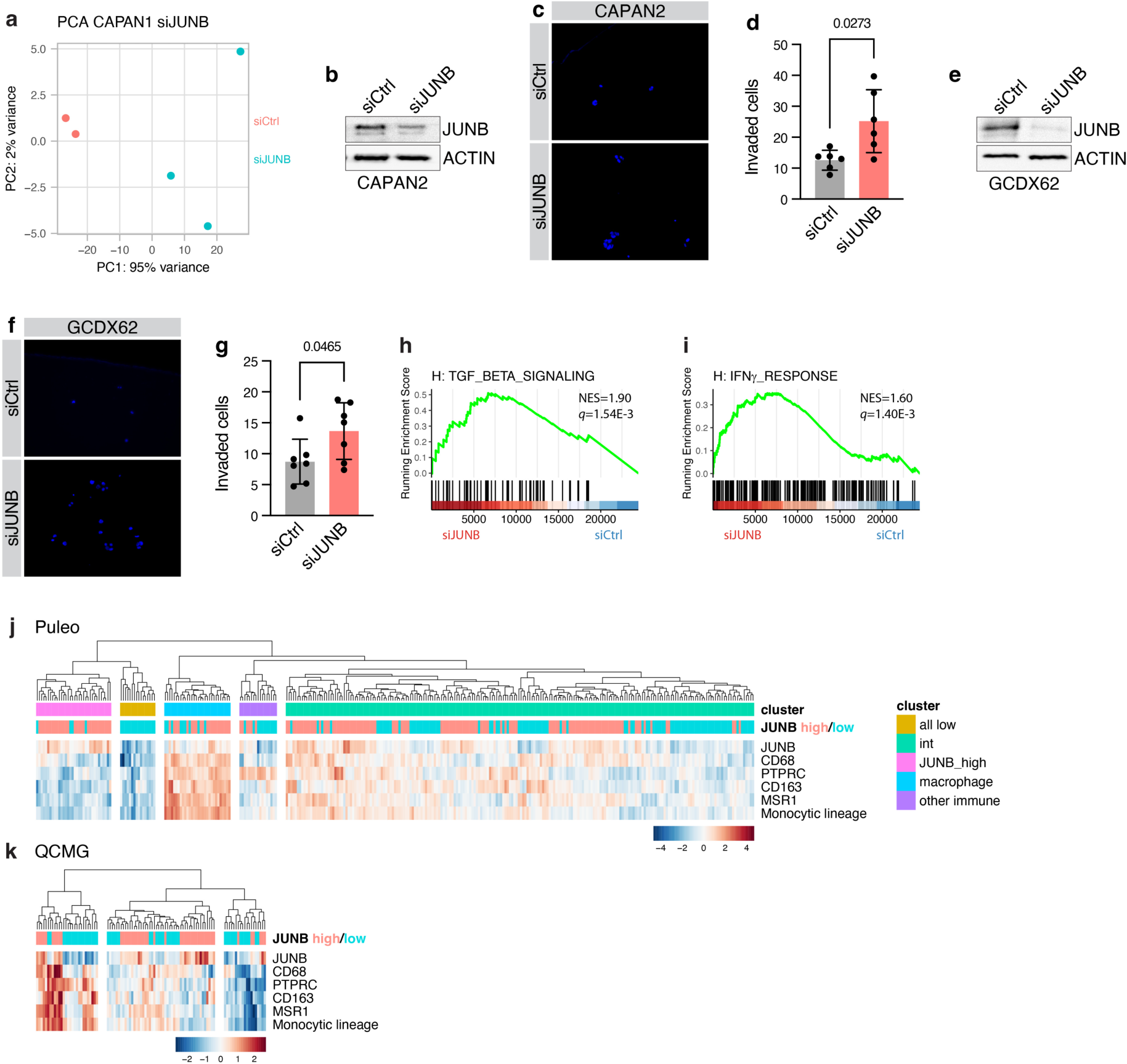
JUNB restricts invasiveness and pro-inflammatory signaling. **a**, PCA plot for RNA-seq data performed for JUNB silencing (siJUNB; n=3) or control siRNA (siCtrl; n=2) in CAPAN1. **b**-**g**, Transwell invasion assay for CAPAN2 (**b**-**d**) and GCDX62 (**e**-**g**) with siJUNB or siCtrl. **b**,**e**, Immunoblot for JUNB and b-actin after siJUNB or siCtrl in CAPAN2 (**b**) and GCDX62 (**e**), validating silencing for the invasion assay. n=3. **c**,**f**, DAPI staining of invaded CAPAN2 (**c**) or GCDX62 (**f**) cells. Scale bar 100 μm. **d**,**g**, Quantification of **c**,**f**, for number of invaded cells. Average counts per FOV with mean ± s.d. shown. **d**, n=6 inserts from n=3 independent experiments. **g**, n=7 inserts from n=4 independent experiments. **h**,**i**, Gene set enrichment analysis plots for ‘TGFβ signaling’ and ‘INFγ response’ Hallmark signatures of the Molecular Signature Database (MSigDB) for siJUNB versus siCtrl in CAPAN1 cells. Normalized enrichment score (NES) and FDR *q* value are indicated. **j**,**k**, Heatmap of Puleo (**j**; n=309) and QCMG (**k**; n=96) expression data for JUNB and macrophage markers as well as MCPcounter scores for the monocytic lineage. JUNB high/low annotation based on top/bottom half of patients for JUNB expression. Cell color indicates z score.

**Extended Data Figure 3.**
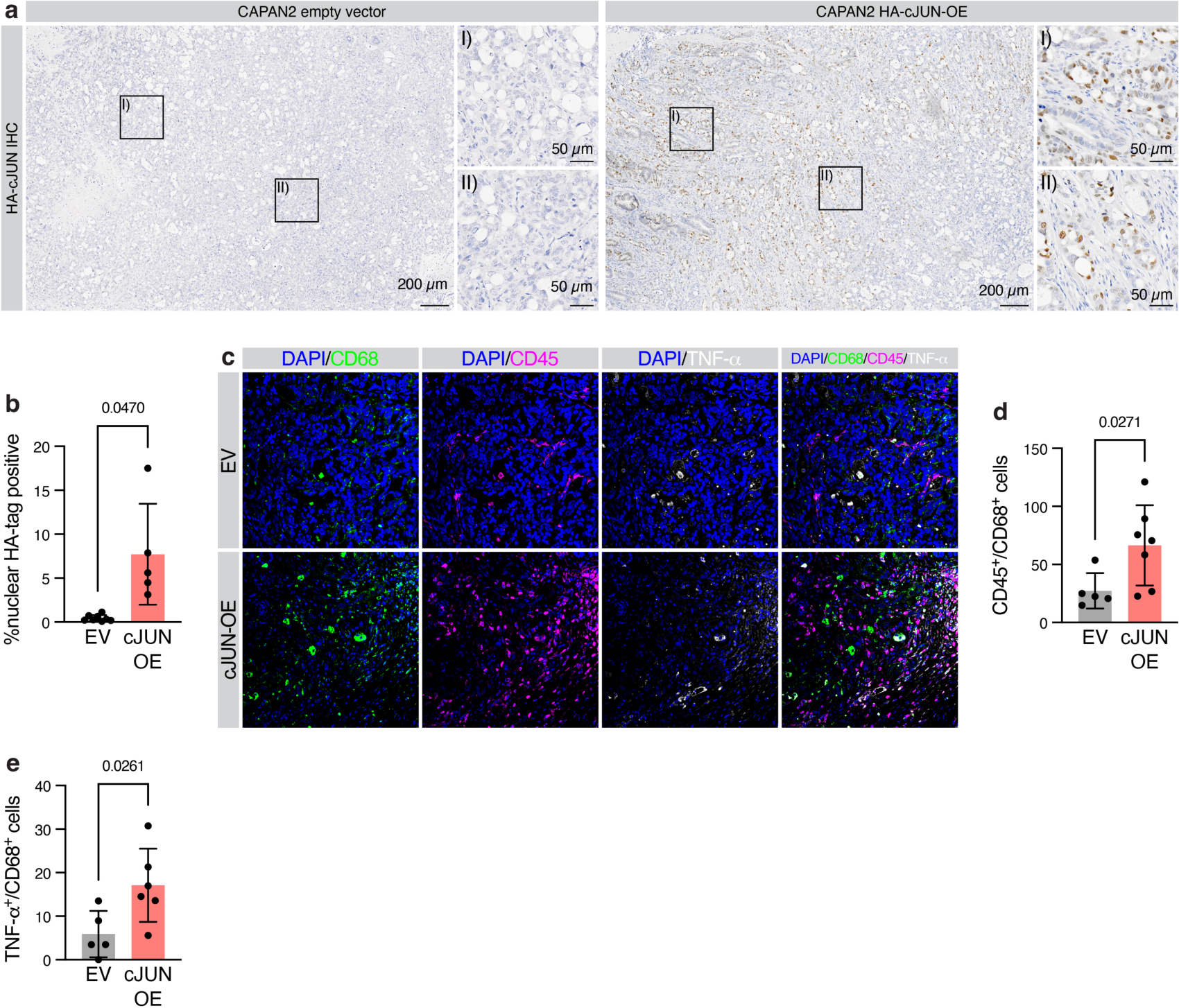
cJUN overexpression enhances TNF-α^+^ macrophage recruitment. **a**-**e**, NMRI-*Foxn1*^nu/nu^ mice were orthotopically transplanted with CAPAN2 cells with stable HA-tagged cJUN-OE or EV control. **a**, IHC for the HA tag of cJUN in orthotopically transplanted CAPAN2 HA-cJUN-OE and EV tumors. Higher magnification insert areas are indicated. Scale bar: overview area, 200 μm; insert, 50 μm. **b**, Quantification of **a** for nuclear HA-tag-positive cells relative to the total number of detected cells with mean ± s.d. shown. **c**, Representative IF staining for CD68, CD45, and TNF-α in orthotopically transplanted CAPAN2 HA-cJUN-OE and EV tumors. Scale bar: 50 μm. **d**,**e**, Quantification of **c** for CD45/CD68 (**d**) and TNF-α/CD68 (**e**) double-positive cells. Per-animal average counts per FOV with mean ± s.d. shown. **d**, EV, n=5; cJUN-OE, n=7. **e**, EV, n=5; cJUN-OE, n=6. **b**,**d**,**e**, Student’s t-test with Welch’s correction.

**Extended Data Figure 4.**
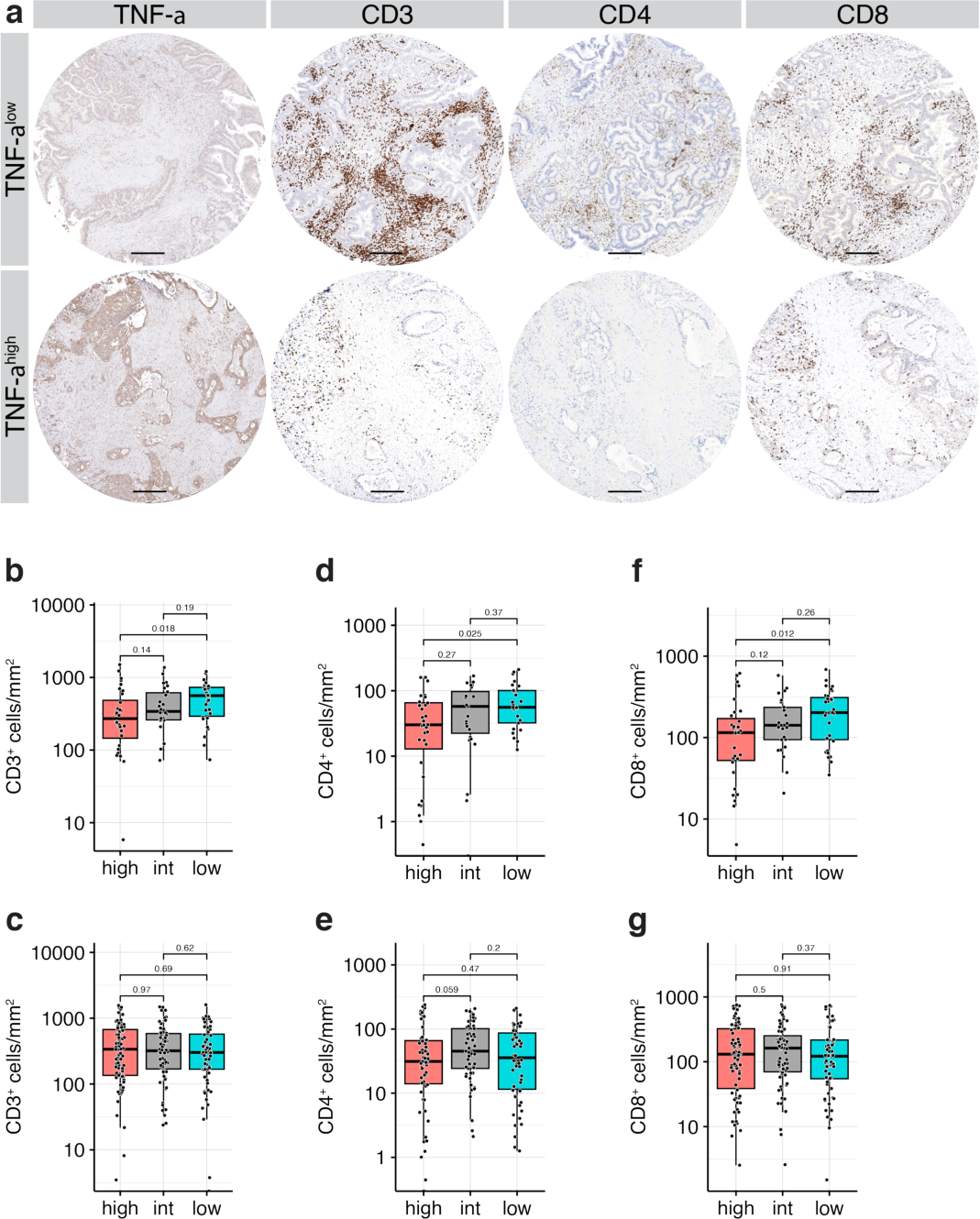
TNF-α expression is associated with T cell exclusion. **a**-**i**, IHC analysis in 105 PDAC patients for TNF-α expression. **a**, IHC for TNF-α, CD3, CD4, and CD8 in cores classified as TNF-α^low^ and TNF-α^high^. Sale bar 200 μm. **b**-**g**, Quantification of **a**, for CD3 (**b**,**c**), CD4 (**d**,**e**), and CD8 (**f**,**g**) in TNF-α^low^, TNF-α^intermediate^ (TNF-α^int^), and TNF-α^high^ expression per patient (**b**,**d**,**f**) and per TMA core across all patients (**c**,**e**,**g**). **h**,**i**, KPC cells were orthotopically implanted into syngeneic C57BL6/J mice and treated with CCR2 inhibitor (CCR2i) RS504393 or vehicle control (VC). **h**, Kaplan-Meier survival analysis. Median survival is indicated. Groups are not significantly different as per log-rank test. **i**, H&E staining of CCR2i and VC tumors. Scale bar 100 µm.

## Notes

### Competing Interest Statement

The authors have declared no competing interest.

